# GRASP: a modular toolkit for building synthetic pentatricopeptide repeat RNA-binding proteins

**DOI:** 10.1101/2025.06.29.661641

**Authors:** Michael Dennis, Su Yi Low, Amy Viljoen, Anuradha Pullakhandam, Catherine Colas des Francs-Small, Leni Campbell-Clause, Charles S. Bond, Ian Small, Farley M. Kwok van der Giezen

## Abstract

Pentatricopeptide repeat (PPR) proteins are eukaryotic RNA binding proteins with multiple roles in mitochondrial and chloroplast transcript processing. PPR proteins are naturally modular and hold great potential for development into tools for RNA processing or controlling RNA folding or expression. However, construction of synthetic PPR proteins is challenging due to their highly repetitive sequences. Here, we present the GRASP kit for assembly of synthetic PPR proteins. Utilising the S-variant of PPR motifs, we designed a library of 42 plasmids which can be combined to assemble synthetic PPR proteins with 9, 14 or 19 motifs to target any RNA sequence of the same length. The GRASP kit enables rapid design and construction of PPR proteins of any desired specificity and is compatible with the MoClo assembly standard. To demonstrate the capabilities of GRASP, we assembled a synthetic PPR RNA editing protein and variants with altered sequence specificity. We tested the functionality of 31 synthetic PPR protein variants against a set of 46 RNA targets and used RNA sequencing to determine levels of RNA editing. The variations in editing provide a wealth of insights into PPR-RNA interactions. The GRASP kit provides a foundation for further development of synthetic PPR protein technologies.

**Graphical abstract:** 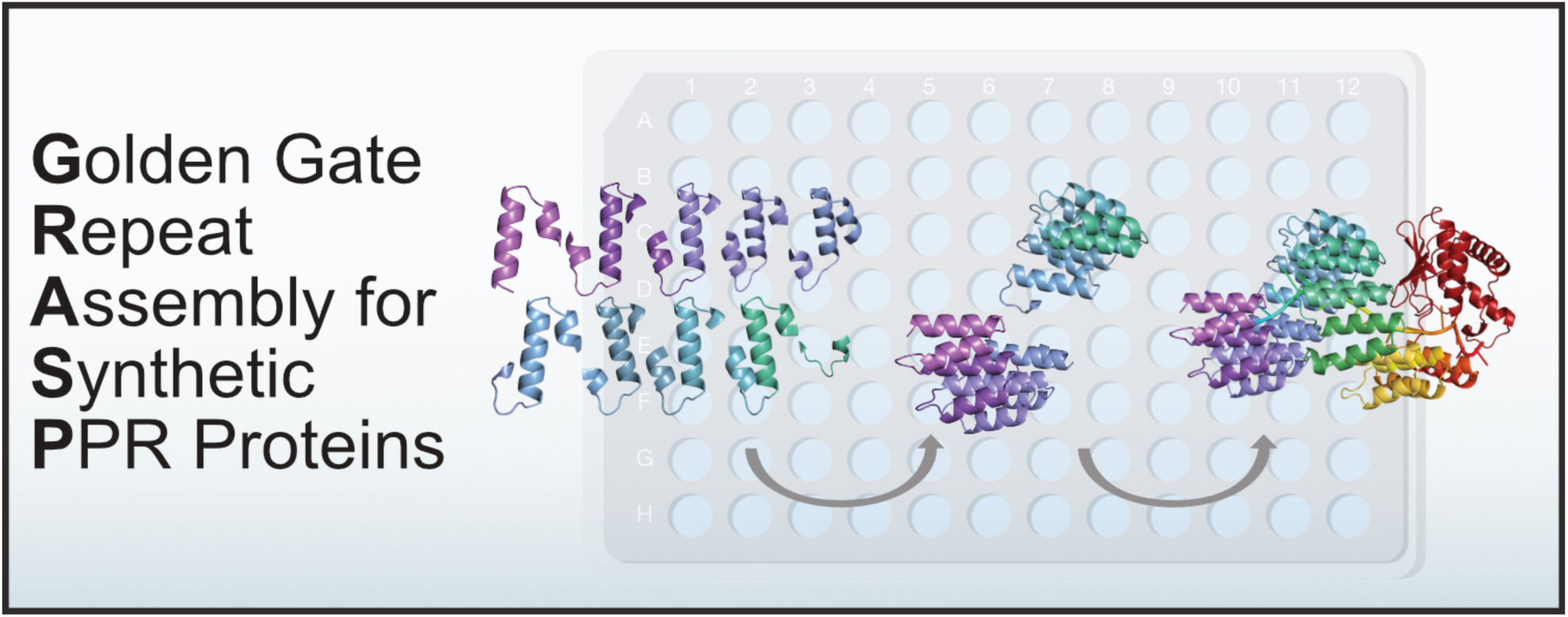

## Introduction

Pentatricopeptide repeat (PPR) proteins are a family of RNA binding proteins present in all eukaryotes, but are particularly numerous in plants (1, 2). They are encoded in the nucleus and predominantly transported to chloroplasts and/or mitochondria where they bind RNA to stabilise (3), cleave (4, 5), participate in splicing (6, 7) or catalyse pyrimidine conversions to edit transcripts (8–10). PPR proteins are ɑ-solenoid proteins composed of a tandem array of 31-36 amino acid helix-turn-helix motifs. They typically encode an N-terminal organelle-targeting peptide, a tandem array of PPR motifs, and in some cases a C-terminal domain for RNA editing (11, 12) or RNA cleavage (5). The tandem array of PPR motifs forms a central binding groove which interacts with RNA in a one-motif, one-nucleotide manner (13). This PPR-RNA interaction has been characterised via a ‘PPR code’ which describes the preferential interactions of PPR motifs with RNA bases (14, 15). The specificity of PPR RNA interactions is largely determined by hydrogen bonds between the fifth and last amino acids of a PPR motif and the aligned RNA base. The PPR code has been used to re-target native PPR proteins to alternate sites (16, 17), and to design novel synthetic PPR proteins to bind RNA (18–20).

Synthetic PPR proteins are attractive for RNA biotechnology due to their naturally modular structure and predictable binding mechanism (21). Native and synthetic PPR proteins have been used to stabilise transcripts (22), and precisely edit RNA both *in vitro* (23) and *in vivo* in plant chloroplasts (20, 24), mitochondria (20, 25) and cytosol (26). They can also function effectively in heterologous systems including bacteria (9, 24, 27, 28) and mammalian cytosol (28, 29). Engineering synthetic PPRs as sequence-specific RNA editing proteins is especially promising due to the high specificity of PPR proteins for their cognate target, their efficiency of converting of cytidine to uridine, and the capacity of some PPR proteins to convert uridine to cytidine (1, 28, 30–33). As PPR specificity is encoded in the peptide sequence, this gives them an advantage for targeting chloroplast and mitochondrial RNA as it avoids the challenge of transporting guide RNAs through organelle membranes that currently inhibits CRISPR-Cas systems (34–36) for this use. Despite this advantage, the repetitive structure of PPR proteins makes them notoriously difficult to manipulate and assemble. Due to their highly repetitive nature, synthetic PPR sequences are often unable to be synthesised via commercial nucleic acid synthesis. These factors have been a barrier for the development of PPR proteins as a tool for manipulating RNA.

A staple of synthetic biology is the Golden Gate assembly method which utilises type IIS restriction enzymes that create a four-base overhang outside of their recognition site (37). Libraries of plasmids compatible with the commonly used MoClo assembly standard (38–41) have been widely adopted, and custom assembly kits have been developed for other modular DNA and RNA binding proteins like TALENs (42–45) and PUFs (46). The naturally modular nature of PPR proteins is well suited to this type of DNA assembly. Golden Gate assembly has been applied to create PPR proteins to interrogate the PPR code (15) and to bind RNA to affect splicing (47), but these methods were designed for bespoke outcomes rather than utility and compatibility with commonly used plasmid libraries (38, 39). Here we present the **<u>G</u>**olden Gate **<u>R</u>**epeat **<u>A</u>**ssembly for **<u>S</u>**ynthetic **<u>P</u>**entatricopeptide repeat proteins (GRASP) kit. GRASP is a MoClo compatible plasmid library for rapid assembly of synthetic PPR proteins to target any RNA sequence. Utilising a consensus motif sequence based on S-type PPR motifs (27, 48), we generated a library of 42 plasmids which can be combined into modules encoding a fully customisable synthetic PPR protein. As a demonstration of the utility of GRASP, we assembled 30 variants of a previously described synthetic PPR protein targeting the *Arabidopsis thaliana* plastid *rpoA* editing site (27), and used a bacterial cell-free expression system to express the proteins alongside 46 variants of the *rpoA* RNA target to profile the effects of mutating individual PPR motifs, and of single nucleotide mismatches in an RNA target. The GRASP kit is a significant step towards optimising synthetic PPR proteins and delivering various PPR protein applications for synthetic biology.

## Materials and Methods

### Design and construction of the GRASP toolkit

The DNA sequence modules of the GRASP kit were based on the “dsnSc” motif sequence of a synthetic pentatricopeptide repeat protein described in (27). GRASP module sequences were designed using Geneious Prime v2021.1. The dsnSc motif was divided and rearranged to accommodate a set of high fidelity four base overhangs compatible with the MoClo Golden Gate assembly standard. GRASP modules act as “CDS1” and “CDS2” modules. Nucleotide sequences for each module were codon randomised using a custom Julia script. GRASP modules were synthesised as gene fragments by Twist Bioscience (San Francisco, California, USA). Gene fragments were suspended in nuclease-free water. The gene fragments were inserted into the universal level -1 vector, pAGM1311 (39) (Supplementary Table S1), and insert sequences were verified by Sanger sequencing performed by Macrogen Inc. (Seoul, South Korea).

### Assembly of 9S_rpoA_DYW using GRASP

GRASP module gene fragments (Supplementary Table S1) were inserted into pAGM9121, the universal level 0 acceptor vector (38). Parts 1A_N, B_DN, C_DT, D_NN, and 1E_NT were used to assemble the CDS1 level 0 construct. Parts 2A_NN, B_NT, C_DN, D_DT, 2E_D were used to assemble the CDS2 level 0 construct. A Golden Gate assembly reaction mix with 0.75 µL BSA (10% w/v), 0.75 µL T4 DNA Ligase Reaction Buffer (New England Biolabs, Ipswich, Massachusetts, USA), 100 U T4 DNA Ligase (New England Biolabs, Ipswich, Massachusetts, USA) and 2.5 U BpiI restriction enzyme (ThermoFisher, Waltham, MA, USA) was added to the plasmid mix, and distilled water was added to a final volume of 10 µL. The reaction was incubated in a thermal cycler set to run 20 seconds at 37°C, followed by 26 cycles of 3 minutes at 37°C and 4 minutes at 16°C, followed by 5 minutes at 50°C, followed by 5 minutes at 80°C. 3µL of the assembly mix was transformed into chemically competent DH5α *E. coli* cells (ThermoFisher, Waltham, Massachusetts, USA). The transformed bacteria were plated onto Lysogeny broth (LB)-Agar with 50 µg/mL spectinomycin, 20 µg/mL X-gal and 0.5 mM IPTG. A starter culture of 5 mL LB media was inoculated with the 50 µg/mL kanamycin. The assembled plasmid was isolated with a FavorPrep Plasmid DNA Extraction Kit (Favorgen Biotech Corp., Pingtung, Taiwan), and the insert sequence was verified by Sanger sequencing performed by Macrogen Inc.

CDS1 and CDS2 plasmids were combined with an N-terminal thioredoxin solubility tag and an RNA editing domain comprised of a triplet of P2-L2-S2 PPR motifs (48) and an E1-E2-DYW catalytic domain previously described in (24). This sPPR coding sequence was placed between a T7 promoter sequence, the *g10-L* ribosome binding site sequence (49) and a modified T7 terminator sequence containing an extended 5ʹ sequence that includes a 16 nucleotide *rpoA* RNA editing site (UUACACGUGCAAAAUC). Level 0 parts were assembled into the pICH47802 level 1 acceptor vector, modified to encode a medium copy origin of replication. A Golden Gate assembly reaction mix was added to the plasmid mix and incubated in a thermal cycler as described above. 3µL of the assembly mix was transformed into chemically competent DH5α *E. coli* cells (ThermoFisher, Waltham, Massachusetts, USA) and the transformed bacteria was plated onto LB Agar with 50 µg/mL carbenicillin, 20 µg/mL X-gal and 0.5mM IPTG. A culture of transformed bacteria was grown at 37°C for 6-8 hours, after which 15 µg/mL chloramphenicol was added to the culture, which was then grown overnight. The level 1 plasmid was isolated using a FavorPrep Plasmid DNA Extraction kit. The insert sequence was verified by Sanger sequencing performed by Macrogen Inc.

### Cell-free expression of 9S_rpoA_DYW

Protein expression of 9S_rpoA_DYW was performed using the NEBexpress cell-free *E. coli* protein synthesis system (New England Biolabs, Ipswich, Massachusetts USA) scaled down to a final volume of 6 µL. A cell-free expression reaction master mix containing 3 µL NEBexpress 2✕ synthesis buffer, 1.44 µL NEBexpress S30 synthesis extract, 6 U T7 RNA polymerase, 5 U Murine RNase inhibitor was combined with 30 ng of the 9S_rpoA_DYW plasmid and made to a final volume of 6 µL with nuclease-free water. The protein expression reaction was incubated at 37°C for 2.5 hours. Following incubation, 0.7 µL of the expression reaction was resuspended in 4 µL 2✕ sample loading buffer (100 mM Tris-HCl, 4% (w/v) sodium dodecyl sulfate (SDS), 20% (v/v) glycerol, 0.2% (w/v) bromophenol blue, 200 mM β-mercaptoethanol) and denatured at 95°C for 5 minutes. All 4.7 µL samples were separated on a 10% Mini-PROTEAN TGX Stain-Free Gel (Bio-Rad Laboratories, Hercules, California, USA) at 120 V for 15 minutes, then 180 V for 40 minutes. Stain-Free Gels were visualised by UV excitation using the Bio-Rad Gel Doc EZ Imaging System with Image Lab software (Bio-Rad Laboratories, Hercules, California, USA). After visualisation, proteins were transferred to an Immun-Blot PVDF membrane (Bio-Rad Laboratories, Hercules, California, USA) using a Trans-Blot SD semi-dry transfer cell (Bio-Rad Laboratories, Hercules, California, USA). The membrane was blocked with 1:10 Western Blocking Reagent (Roche Diagnostics GmbH, Mannheim, Germany) for 1 hour at room temperature. After blocking, the membrane was rinsed briefly in Tris Buffered Saline with 0.1% Tween 20 (TBST) and then incubated with primary 6✕-His Tag Mouse Monoclonal Antibody (ThermoFisher, Waltham, Massachusetts, USA) diluted to 1:4000, at 4°C overnight or at room temperature for 3 hours. The membrane was washed with TBST three times for 5 minutes each and incubated with a rabbit anti-mouse secondary antibody conjugated to horseradish peroxidase (ThermoFisher, Waltham, Massachusetts, USA), diluted to 1:10 000, at room temperature for 1 hour. The membrane was then washed five times for 5 minutes each in TBST and incubated with SuperSignal West Pico PLUS Chemiluminescent Substrate (ThermoFisher, Waltham, Massachusetts, USA) for 1 minute. Visualization was done using an Amersham Imager 680 (GE Life Sciences Chicago, Illinois, USA).

Following incubation, 2 µL of 10✕ RQ1 DNase Buffer, 6 µL of 50% PEG4000, and 1U RQ1 DNase (Promega, Madison, Wisconsin, USA) was added to the expression reaction, and nuclease-free water was added to make a final volume of 20 µL. Plasmid DNA was digested at 37°C for 30 minutes. 1 µL (20 µg) RNA Grade Proteinase K (ThermoFisher, Waltham, Massachusetts, USA) was added to the supernatant and incubated at 65°C for 20 minutes. 3 µL of the treated expression reaction was mixed with 1 µL of 10 µM reverse transcription primer, and Proteinase K was inactivated by incubation at 80°C for 10 minutes. The Proteinase-K-treated reaction was held at 4°C for 3 minutes and then reverse-transcribed by ProtoScript II (New England Biolabs Ipswich, Massachusetts, USA) according to the manufacturer’s instructions. A 2 µL aliquot of cDNA was PCR amplified using recombinant *Taq* DNA Polymerase (ThermoFisher, Waltham, Massachusetts, USA). The PCR product was sequenced by Macrogen Inc. Sanger chromatograms were trimmed and RNA editing was quantified using MultiEditR (50).

### Assembly of 9S_rpoA_DYW protein variants

GRASP plasmids were used to assemble a set of 31 sPPR variants of 9S_rpoA_DYW with systematic modifications at each S-motif and the P2-motif to target each alternative RNA base (Supplementary Table S2). Golden Gate assembly and molecular cloning of GRASP level 0 CDS plasmids and level 1 constructs was performed as described earlier in the Methods for assembly of 9S_rpoA_DYW using GRASP, except with an unmodified T7 terminator part. The 31 9S_rpoA_DYW protein variant insert sequences were verified by Sanger sequencing performed by Macrogen Inc.

### *In vitro* transcription of RNA targets for a high-throughput editing assay

RNA targets were designed in Geneious Prime v2023.1 and synthesised as DNA oligonucleotides by Twist Biosciences. The oligonucleotides included a T7 Promoter for *in vitro* transcription and encoded the 16-nucleotide *rpoA* target of 9S_rpoA_DYW, or a variant where each nucleotide except the edit site was systematically altered. Oligonucleotides encoded ambiguities B (A → C/G/T), D (C → A/G/T), H (G → A/C/T), or V (T → A/C/G) in each position of the *rpoA* target, resulting in a set of 46 variants. The *rpoA* targets were flanked by primer binding sites, and 5ʹ and 3ʹ stem-loop sequences (51, 52) (Supplementary Table S3).

Lyophilised oligonucleotides were resuspended in nuclease-free water to a final concentration of 10 ng/µL and pooled to make an equimolar mix. In triplicate, 100 µL of pooled oligonucleotides (1 µg of DNA) were dried and resuspended in 10 µL nuclease-free water. The DNA oligonucleotides were transcribed into RNA using T7 RNA Polymerase (New England Biolabs, Ipswich, Massachusetts, USA) in an *in vitro* transcription mix containing RNase Inhibitor (New England Biolabs, Ipswich, Massachusetts, USA) and 5 mM DTT, and incubated overnight at 37-°C. Triplicates of the *in vitro* transcription mix were pooled and the DNA template was digested with 1 U RQ1 DNase in a final volume of 100 µL. Transcribed RNA was mixed with 1 volume of pH 4.7 phenol:chloroform (5:1), vortexed for 1 minute, and centrifuged for 2 minutes at 12000 *g*. The aqueous phase was isolated and mixed with 1 volume of chloroform:isoamyl (24:1), vortexed, and centrifuged for 2 minutes at 12000 *g*. The aqueous phase was recovered and 0.5 volumes of 7.5M ammonium acetate and 2.5 volumes of 100% ethanol were added. RNA was incubated at -80°C for 30 minutes, and then centrifuged at 18000 *g* for 20 minutes at 4°C. The RNA pellet was washed with 1 mL of 70% ethanol and then centrifuged at 18000 *g* for 20 minutes at 4°C. The RNA pellet was dried at room temperature and resuspended in 20 µL nuclease-free water. The final concentration of the RNA target pool was determined by Qubit HS RNA (Invitrogen, Waltham, USA).

### High-throughput cell-free protein expression

Cell-free protein expression of 9S_rpoA_DYW and 30 9S_rpoA_DYW variants was performed in triplicate as described earlier in the Methods for cell-free expression of 9S_rpoA_DYW. 0.5 µL of a DMSO solution containing 325 µM chloramphenicol and 13✕ proteinase inhibitor cocktail for His-tagged proteins (Sigma Aldrich, St. Louis, MO, USA) was added to inhibit further protein expression. A 1µL aliquot of each reaction was mixed with 39 µL of Laemmli buffer, denatured at 95°C, snap-frozen, and stored at -80°C. A 1 µL aliquot of the expression reactions was diluted 1:20 in 1✕ *in vitro* Editing Buffer (30 mM HEPES-KOH pH 7.7, 3 mM magnesium acetate, 45 mM potassium acetate, 30 mM ammonium acetate, 2 mM ATP, 5 mM DTT, 1% PEG 6000, 5% glycerol). An aliquot of the 46-target RNA pool was diluted to 780 pM, wherein each target is expected to be at ∼17 pM. This was denatured at 80°C for 3 minutes, then cooled to 4°C. To each reaction, 0.32 µL of the RNA pool was added, resulting in a total amount of 250 amol (∼5.4 amol of each target) per reaction. The protein and RNA reaction mix was incubated at 37°C for 2.5 hours. Each reaction was treated with 1 U RQ1 DNase treatment in a final volume of 10 µL. Reactions were treated with proteinase K as described previously in the Methods for cell-free expression of 9S_rpoA_DYW. A 3 µL aliquot of each reaction was mixed with 1 µL of 10 µM NGS_RT_Generic primer (Supplementary Table S4). Reverse transcription was performed as described earlier in the Methods for cell-free expression of 9S_rpoA_DYW. Custom-indexed PCR primers (Supplementary Table S4) were added into a 96-well plate with 2 µL of cDNA. Each reaction was amplified using a recombinant *Taq* DNA Polymerase master mix (5 µL 10x *Taq* Buffer, 5 µL 2mM dNTPs, 5 µL 25 mM MgCl2, 34.75 µL water, 0.25 µL *Taq* DNA Polymerase) (ThermoFisher, Waltham, Massachusetts, USA). The cDNA products were amplified with the following program: 30 seconds at 75°C for 5 cycles, 30 seconds at 72°C with a decreasing ramp of 1°C per cycle for 8 cycles, 10 seconds at 72°C, and 30 seconds at 64°C for 15 cycles, 10 seconds at 72°C.

### RNA sequencing

cDNA from the cell-free expression reactions was pooled and vacuum-dried, resuspended in 50 µL distilled water, and cleaned by mixing with 150 µL of TE Buffer (pH 8.0), 100 µL of 30% PEG 8000, 30 mM MgCl_2_ and 1 µL linear acrylamide (ThermoFisher, Waltham, MA, USA). After vigorous vortexing, a DNA pellet was recovered by centrifugation at room temperature for >20000 *ɡ* for 30 minutes. The pellet was resuspended in distilled water and run on an 8% TBE acrylamide gel. Bands corresponding to 71 bp were excised and placed in a 0.5 mL microcentrifuge tube, punctured at the base. The 0.5 mL microcentrifuge tube was placed in a 2 mL microcentrifuge tube and centrifuged at 10000 *ɡ* for 5 minutes. Extruded acrylamide slurry was mixed with 250 µL of TE Buffer and incubated for 2 hours at 25°C on a shaking thermomixer. The TE buffer and acrylamide slurry were centrifuged through a Spin-X Cellulose Acetate Column (Corning Inc., Corning, NY, USA). The flow through was mixed with 1/10 volume of 3 M sodium acetate, pH 5.2, 3 volumes of ice-cold ethanol, and 1 µL of linear acrylamide. cDNA was precipitated at -80°C for 30 minutes and then centrifuged at 18000 *ɡ* for 30 minutes at 4°C. The cDNA pellet was washed with 1 mL of 80% ethanol, and centrifugation again at 4°C at 18000 *ɡ* for 30 minutes. The cDNA Pellet was dried at room temperature and resuspended in 20 µL distilled water. A 12.5 µL aliquot used to generate a DNA sequencing library using a TruSeq DNA Nano kit (Illumina, San Diego, California, USA). The cDNA pools was 3ʹ adenylated, and adaptors were ligated according to the manufacturer’s protocol. The adaptor-ligated pool was purified using a QiaQuick PCR Purification kit (Qiagen, Hilden, Germany). The purified amplicon library was eluted in 50 µL of distilled water. A 25 µL aliquot of the eluate was enriched by 8 cycles of PCR, and purified using a QiaQuick PCR Purification kit. The purified library was run on a TBE 8% acrylamide gel and a band corresponding to 192 bp was excised and purified using a Spin-X Column as before. The amplicon library was quantified by qPCR using a KAPA Library Quantification Kit for Illumina Platforms (Roche, Basel, Switzerland). Sequencing was performed on an Illumina MiSeq set to 75 reads in paired-end mode using a MiSeq Reagent Kit v3 (Illumina, San Diego, California, USA).

### RNA editing detection and analysis

RNAseq reads were merged using bbmerge (53) with default settings. Approximately 95% of read pairs could be merged successfully to reconstruct the expected 71 nt insert. Each read in each library was assigned to one of the 96 reactions using the row and column index barcodes within the insert. The edit site was identified using the immediately adjacent upstream sequence (TTACACGTGCAAAAT) as a query (allowing mismatches, as implemented by ApproximateSearchQuery in BioSequences.jl (github.com/BioJulia/BioSequences.jl). Edited and unedited reads were counted for each target variant and each well. The number of reads per variant in each well ranged from 200–9000. The counts were converted to proportions; the proportion of edited reads varied from 0.0 to 0.23 in the experiments shown. The proportions were used to train a logistic regression model using GLM.jl (formula: proportion of edited reads ∼ RNA target + protein variant + interaction between RNA target & protein variant; binomial distribution; logit link) (github.com/JuliaStats/GLM.jl). This model thus has 30 coefficients for the protein variants, 45 for the RNA variants and 1350 coefficients for the interactions between them. For analyses involving combining effects observed in multiple protein/RNA combinations, the set of comparable regression coefficients was collected and the maximum absolute value taken as the representative value; this is referred to as the maximum pooled coefficient.

### Analysis of sequences of natural PPR binding sites and natural PPR proteins

Natural PPR binding sites in chloroplasts from *Selaginella kraussiana*, *Selaginella lepidophylla* and *Selaginella uncinata* were extracted using data from (54, 55). The reported editing sites were first used to edit the genome sequence (C to T transitions at the reported sites), then the sequence from -15 to -6 with respect to each reported site was extracted and used to analyse base composition. An identical analysis was done using data for angiosperm mitochondria and chloroplasts (see github repository listed in the data availability statement for data sources). The sequences of natural S-class PPR proteins were obtained from our previous analysis (1) of the 1KP transcriptome dataset (56). ‘SS’ motif sequences (as defined by (48)) were extracted and filtered to retain only those that were 31 amino acids long with the 5th and 31st amino acid pair being one of ND, NN, TD or TN. This constituted a set of 47878 motifs that were used to study the relationship between the net charge of the motif (calculated at pH 7.7) and the 5th/31st combination. This set of motifs was also used to investigate how the amino acid preference at position 29 differed between motifs with N or D at position 31. Preference was calculated as the mean difference in probability, where the probability density functions were assumed to be beta distributions, Beta(α_n_ + 1, β_n_ + 1) where α_n_ = count of motifs with a particular amino acid at position n and β_n_ = count of motifs with other amino acids at position n.

## Results

### An assembly toolkit for synthetic pentatricopeptide repeat proteins

The GRASP kit is a set of MoClo-compatible vectors for type IIS (Golden Gate) assembly of synthetic PPR (sPPR) proteins based on the consensus “dsnSc” S-type motif described in (27) (Fig. 1A). Typically, PPR RNA editing proteins consist of repeating triplets of P1-(35 aa) L1-(35 aa) and S1 (31 aa) motifs (48), followed by an RNA editing domain consisting of a triplet of P2 (35 aa) L2 (36 aa) S2 (32 aa) PPR motifs, and a cytidine deaminase-like E1-E2-DYW domain (8, 48). Some plants encode PPR RNA editing proteins with an alternative motif arrangement consisting of repeats of a 31 amino acid “S” motif (formerly “SS”), followed by the RNA editing domain (1). This arrangement resembles P-class PPR proteins, which consist of a tandem array of 35 amino acid P-motifs. P-class PPR proteins are predominantly involved in transcript stabilisation (57–59), but can also contain C-terminal domains implicated in RNA cleavage (4, 5). S-type PPR RNA editing factors can independently catalyse the conversion of cytidine to uridine in RNA transcripts (27), and their single motif PPR array architecture allowed for the design of a compact set of 24 plasmids to assemble a 9 motif sPPR protein that can target any 9 nucleotide RNA sequence.

**Figure 1.**
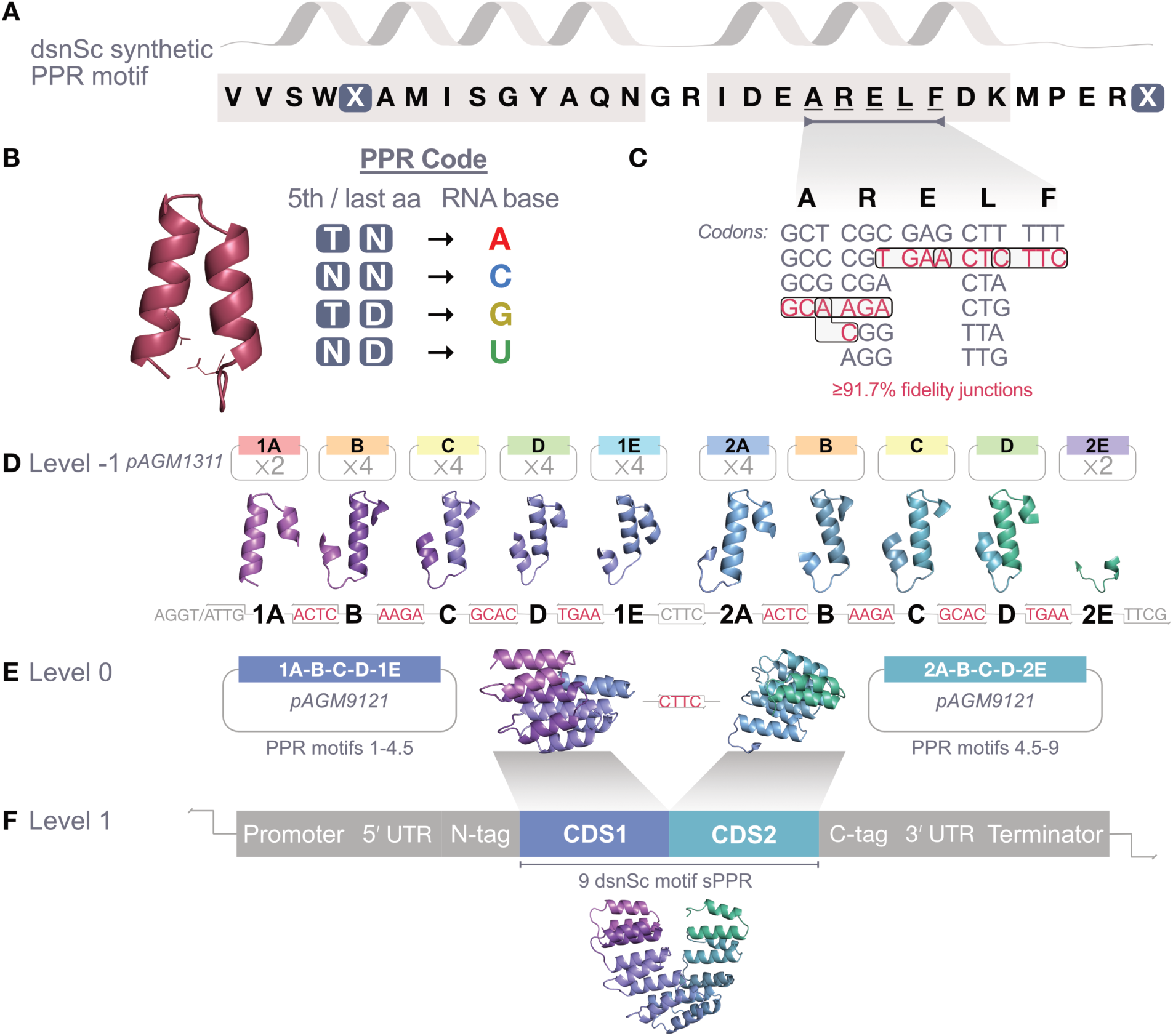
Schematic of the GRASP kit for assembly of a 9-motif synthetic pentatricopeptide repeat protein. (**A**) GRASP uses the “dsnSc” 31 amino acid S-type PPR motif described in (27). (**B**) Grasp utilises a minimal motif-to-base preference code of Thr/Asn for binding adenine, Asn/Asn for binding cytidine, Thr/Asp for binding guanine and Asn/Asp for binding uridine (14, 15). (**C**) The region between amino acids 20-24 (ARELF) contains sufficient codon diversity to allow for selection of high-fidelity four-base Golden Gate overhangs from those experimentally profiled in (60). (**D**) Modules in the GRASP kit are encoded on the universal level -1 acceptor vector pAGM1311 (39). Each module codes for one or two partial dsnSc motifs and are labelled alphabetically A through E. Each module comes in four variants, each with variants of the last and 5th residues of amino acids involved in specifically targeting A, C, G or U. An additional 16 modules for assembly of 14 and 19 motif sPPR proteins are included in the GRASP kit (Supplementary Table S1) (**E**) 1A, B, C, D and 1E, and 2A, B, C, D and 2E modules can be combined in separate reactions to assemble two GRASP CDS modules inserted into the universal level 0 acceptor vector pAGM9121 (38). Each CDS module encodes 4.5 dsnSc motif sequences. (**F**) GRASP CDS1 and CDS2 act as coding sequence modules in the MoClo (38) Golden Gate assembly standard. When assembled into a level 1 transcriptional unit, GRASP CDS1 and CDS2 modules combine to encode a 9-motif sPPR protein. The sPPR protein can be combined with a variety of optional N-or C-terminal protein domains.

GRASP utilises a minimal PPR code of four strong interactions between the fifth and last amino acids of a dsnSc motif and an RNA base (Fig. 1B) (14, 15, 27). Due to the critical importance of the 5th and last amino acid in the dsnSc motif for RNA base recognition, and lack of codon diversity at position 1 and 31 in the dsnSc motif, assembly junction points for dsnSc motifs were placed at a region of high codon diversity between amino acids 20-24. Codon diversity in this region allowed for codon optimisation of GRASP modules to encode a set of high-fidelity four base overhangs compatible with the MoClo molecular cloning standard (Fig. 1C) (60).

Using the GRASP kit, a 9-motif sPPR protein is assembled in two stages. Twenty-four GRASP modules encoded in the universal level -1 vector pAGM1311 (39) are named numerically 1 and 2, and alphabetically A, B, C, D and E, according to the order of module assembly (Fig. 1D). “A” modules encode four-base assembly junction points that match “B” modules, “B” modules match “C” modules, etc. In the first stage of assembly, two separate level 0 constructs containing parts 1A-B-C-D-1E and 2A-B-C-D-2E are inserted into the universal level 0 acceptor vector pAGM9121 (38). The resulting coding sequences (CDS) encode 4.5 dsnSc motifs with 5th and last amino acids for RNA base specificity varying depending on the GRASP modules selected (Fig. 1E). In the second stage of assembly, these CDS modules are combined to form the 9 dsnSc motif RNA binding tract in a level 1 transcriptional unit assembly with the user’s selection of promoter, UTR, effector domain and terminator elements (Fig. 1F). Optionally, an additional 14 level -1 GRASP modules were designed to extend sPPR proteins by 5 or 10 motifs. These can be used to assemble 14 or 19 motif dsnSc RNA binding tracts (Supplementary Fig. S1 and Supplementary Table S1).

### Reconstruction of TRX-9S-DYW sPPR RNA editing factor

Several recent studies of synthetic PPR RNA editing factors have utilised the *Arabidopsis thaliana* plastid *rpoA* target of the editing factor CHLOROPLAST BIOGENESIS 19 (CLB19) (24, 27, 28, 61). Synthetic PPR RNA editing factors targeting the *rpoA* site edit efficiently in both *E. coli*, and *in planta* (24, 27). To test the efficacy of the GRASP kit, we reconstructed the synthetic protein “TRX-9S-DYW” from (27) which targets the CLB19 *rpoA* sequence (27). TRX-9S-DYW utilised four variants of the S-type motif, whereas the GRASP synthetic reconstruction (hereafter named 9S_rpoA_DYW) utilises only the dsnSc consensus S-motif sequence (Fig. 1A). 9S_rpoA_DYW was assembled with an N-terminal thioredoxin solubility tag, and a triplet of P2-L2-S2 PPR motifs that link the GRASP dsnSc motifs to a C-terminal E1-E2-DYW RNA editing domain, both previously described in (24). The 9S_rpoA_DYW transcript is under the control of the T7 promoter, and a modified T7 terminator module which includes 33 bases upstream and 5 bases downstream of the *rpoA* RNA editing site (Fig. 2A). Transcription of the 9S_rpoA_DYW construct runs through the *rpoA* target RNA editing site, thus 9S_rpoA_DYW edits the same transcript it is translated from (Fig. 2A). 9S_rpoA_DYW was expressed in bacterial cell-free expression lysate (Fig. 2B) and edited the co-transcribed *rpoA* target at an average efficiency of 38.69% (Fig. 2C). A 19 motif sPPR protein (19S_rpoA_DYW) targeting an extended sequence of the *rpoA* binding site was also assembled and assayed and was capable of RNA editing up to 49.28% (Supplementary Fig. S2).

**Figure 2.**
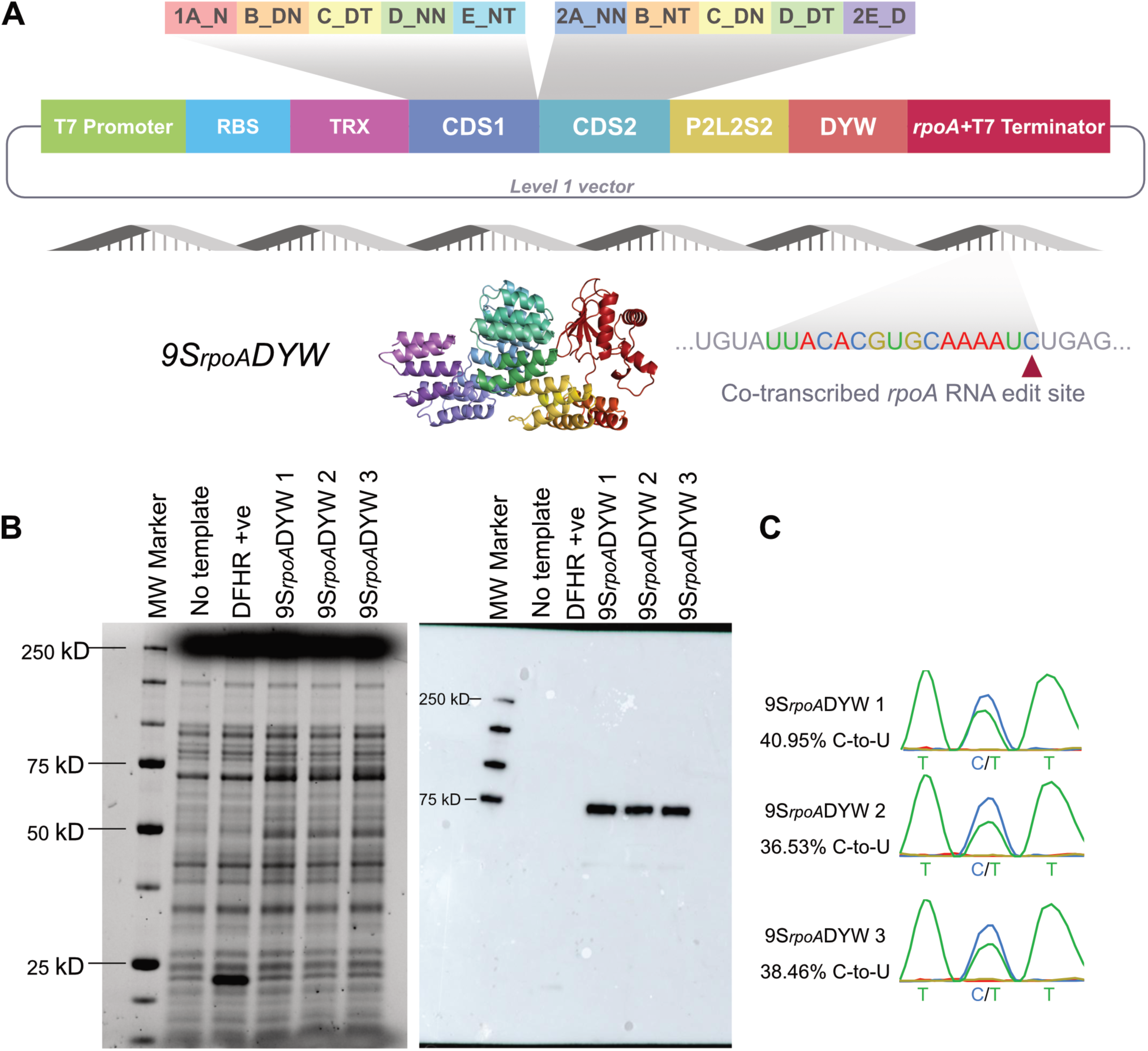
Reconstruction and functional assay of 9SrpoADYW. (**A**) The TRX-9S-DYW synthetic PPR protein (27) was reconstructed using GRASP modules, with an N-terminal thioredoxin solubility tag, and a C-terminal consensus RNA editing domain (24). We named this iteration 9SrpoADYW. The *rpoA* binding and editing site is positioned after the 9SrpoADYW stop codon, so 9SrpoADYW edits the same transcript it is translated from. (**B**) SDS-PAGE showing expression of 9SrpoADYW in an *E. coli* cell-free expression reaction and Western blot showing 9SrpoADYW detected using anti-His tag antibody. Full acrylamide gel and western blot images are provided in Supplementary Fig. S3. (**C**) Sanger chromatograms of cDNA amplified from *E. coli* cell-free lysate expression reactions shows that 9SrpoADYW assembled with the GRASP kit is functional. Sanger chromatogram peak proportions were determined using MultiEditR (50).

### High-throughput assembly and activity assay of synthetic RNA editing proteins

Next, to demonstrate the utility of the GRASP kit for high-throughput assembly of sPPR proteins and to interrogate the specificity of 9S_rpoA_DYW, we designed 30 variants of 9S_rpoA_DYW (p1-p30). Variants p1 to p30 encode the 9S_rpoA_DYW sPPR protein, with systematic modification of each dsnSc motif, and the P2 motif of the P2-L2-S2 triplet. These ten modified motifs are hereafter referred to as m1 to m10. Motifs were modified such that each targets an alternative RNA base relative to 9S_rpoA_DYW. For example, m1 of 9S_rpoA_DYW encodes the GRASP 5th-last amino acid combination of asparagine (N) and aspartate (D) which we expected to target uridine. Variants p1 to p3 encode the GRASP 5th-last amino acid combination for targeting adenine (TN), cytidine (NN) and guanine (TD), and so on for motifs m2-m10 (Fig. 3A). We expected each motif variant to show differing sequence specificity at the associated RNA base.

**Figure 3.**
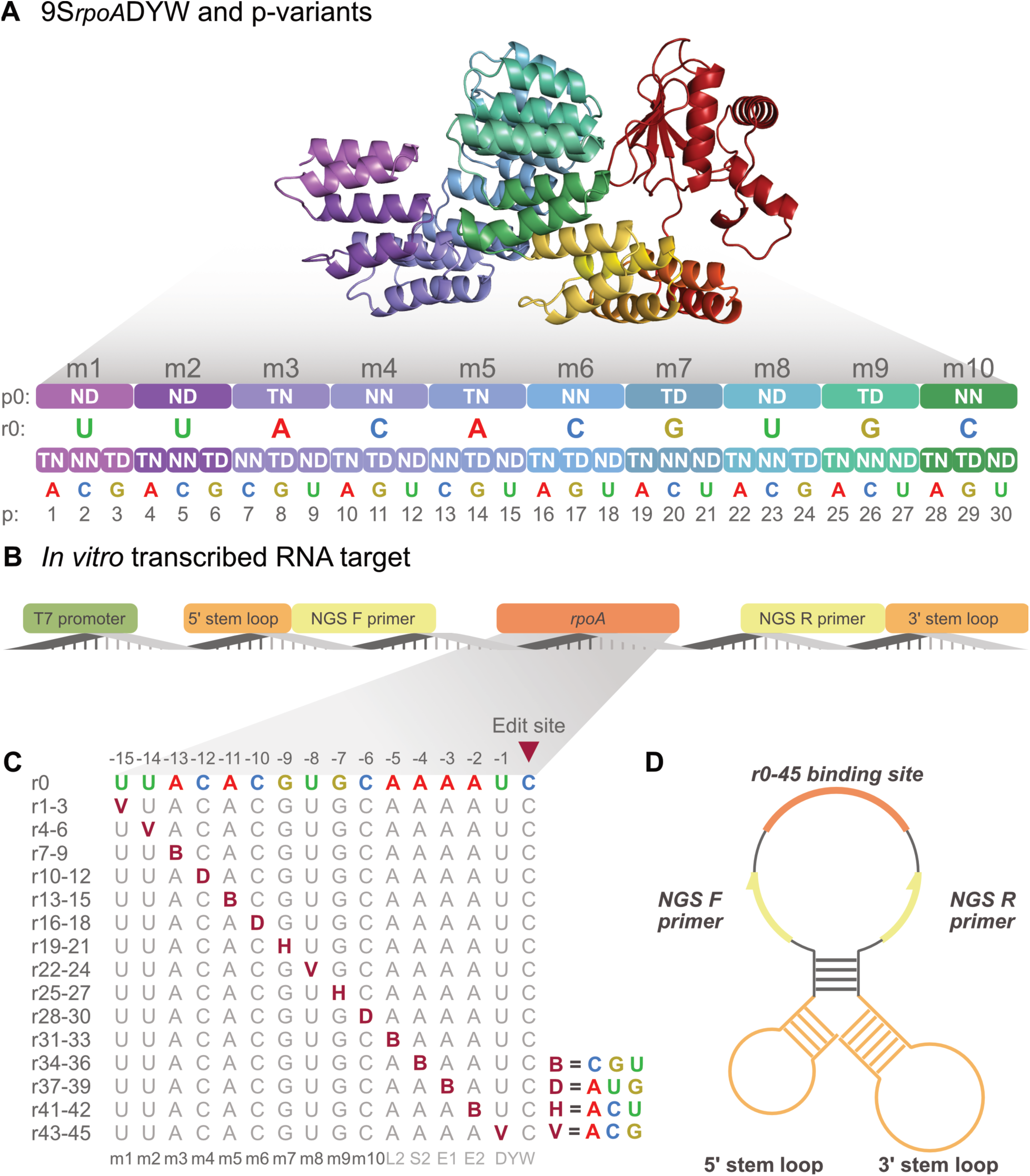
9SrpoADYW ‘p-variants’ and *in vitro* transcribed 9SrpoADYW RNA targets and ‘r-variants’. (**A**) AlphaFold 3 (63) model showing 9SrpoADYW with RNA base preferences for dsnSc motifs (m1-m9) and the P2 motif (m10), and the ‘p-variants’ with altered RNA base preference. Motif colours in the lower part of the panel match the colours in the structural model. Protein model visualisation made in PyMol (64). (**B**) Synthesised gene fragments encoding a 129 nucleotide RNA target under the T7 promoter. The original *rpoA* target of 9SrpoADYW and 45 sequence variants are flanked by custom primer binding sites for preparation of NGS sequence libraries, and a 5ʹ and 3ʹ stem loop sequence (51, 52) to stabilise the transcript after *in vitro* transcription. (**C**) RNA variants of the *rpoA* sequence (r2-45) were modified at positions 15 to 1 (upstream relative to the edited C) with ambiguities representing each alternate RNA base. The resulting 45 targets were pooled in equimolar concentrations and included in each 9SrpoADYW and variant expression reaction. Alignment of the dsnSc motifs and the RNA editing domain (P2-L2-S2-E1-E2-DYW) are shown below. (**D**) After *in vitro* transcription, the 129 nucleotide RNA scaffold is predicted to fold into an open loop flanked by the 5ʹ and 3ʹ stem loop structures. Secondary structure predictions were performed using Mfold (Supplementary Table S5) (65).

To control the concentration of RNA accessible for editing by 9S_rpoA_DYW and p1-p30, we decoupled the RNA editing target from the protein coding transcript. We designed a 129 nucleotide RNA sequence that includes 5ʹ and 3ʹ stem loop structures (51, 52) for transcript stabilisation, and primer binding sites for target amplification flanking the *rpoA* RNA editing site (r0) (Fig. 3C). The RNA scaffold was placed under the T7 promoter for *in vitro* transcription. Secondary structure predictions of the RNA scaffold predict an accessible open loop stabilised by the flanking stem loop structures (Fig. 3D) (62). We synthesised 45 variants of the *rpoA* RNA target (r1-r45) such that in each of the individual 15 positions upstream of the cytidine editing target, every RNA base was represented (Fig. 3C). Equimolar r variants were pooled and transcribed, then included in each p variant cell-free expression reaction, therefore every r-variant was available to be edited by every p-variant. The RNA pool from each reaction was reverse transcribed, and the cDNA was used to prepare a 71 nt stranded sequencing library which was sequenced on an Illumina MiSeq. The 46 r-variants (r0-r45) were edited between 0%-23% by the 31 p-variants (Fig. 4A).

**Figure 4.**
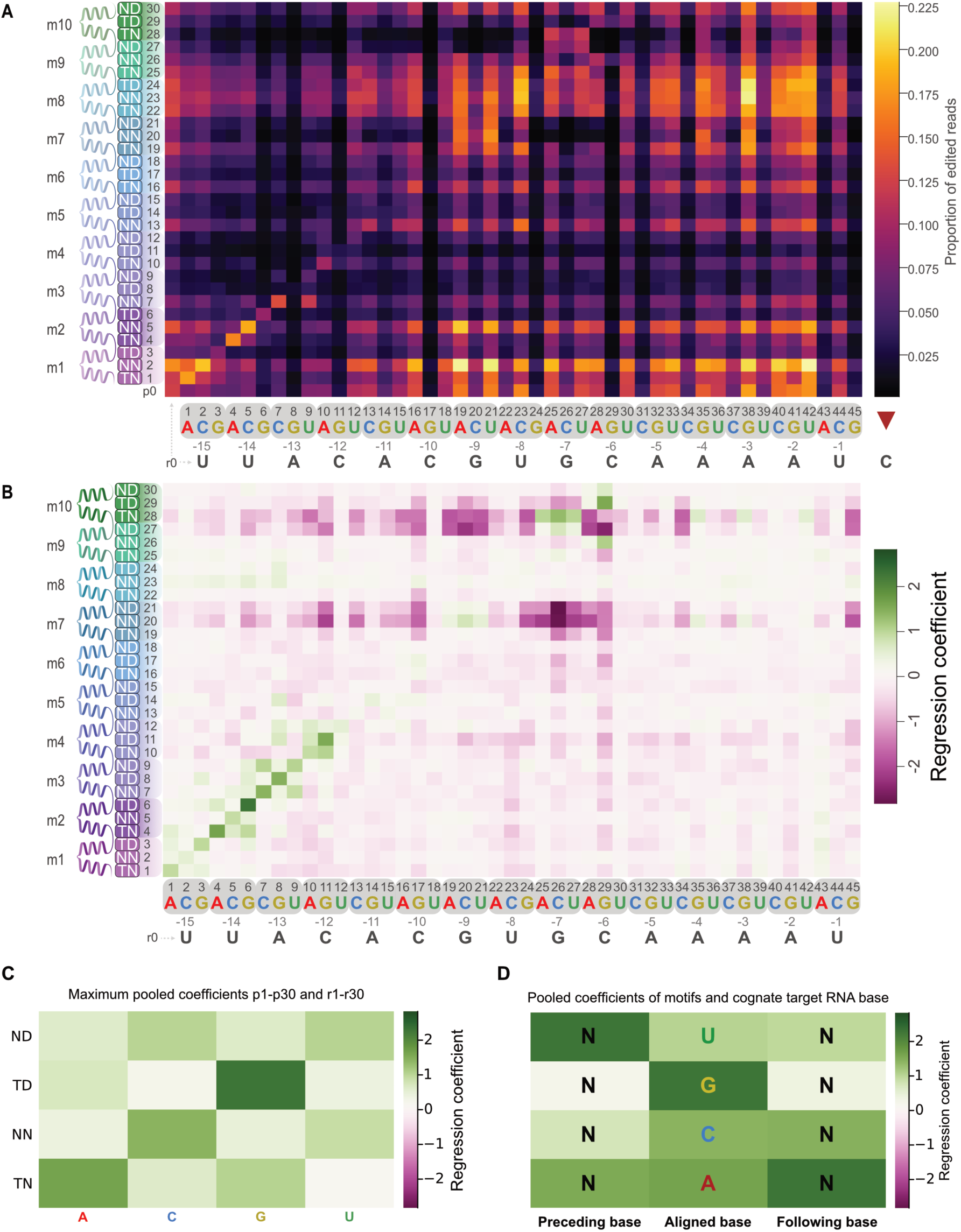
Heatmaps displaying editing of RNA targets catalysed by 9SrpoADYW and its variants, and model coefficients from a logistic regression analysis of the RNA editing data. In all cases the reference is the editing seen with the unmodified protein (p0) and/or the unmodified target (r0). (**A**) 9SrpoADYW (p0) and 30 modified variants (p1-p30) were expressed in *E. coli* cell-free expression lysate and incubated with a pool of *in vitro* transcribed RNA targets comprising *rpoA* (r0) and single nucleotide sequence variants (r1-45). The values indicate the mean (n = 3) proportion of edited reads; (**B**) Protein:RNA interaction coefficients indicating ‘specificity’; (**C**) Maximum pooled interaction coefficients for cognate interactions between p1:r1 to p30:r30 shows the base preference for each dsnSc motif variant; (**D**) Maximum pooled interaction coefficients preceding and following cognate motif/base interactions.

### Synthetic RNA editing proteins preferentially edit their cognate RNA targets

Thirty-one variations of the 9S_rpoA_DYW synthetic RNA editing factor (p0-p30) were expressed and incubated with 46 variants of the *rpoA* target RNA editing site (r0-r45) resulting in 1426 combinations per replicate (n=3) (Fig. 4A). For each p variant, a corresponding r variant with an optimal match according to the GRASP PPR code was present in the reaction. We expected these optimal pairs to exhibit higher editing efficiency over the 45 additional ‘off-target’ single nucleotide variants of the RNA binding site. We performed a logistic regression analysis of the data, assuming that the observed proportions of edited transcripts were a function of the editability of the RNA target, the activity of the protein and the strength of the specific protein:RNA interaction (Fig. 4B, Fig. 5A, Fig. 6A). Plotting the protein:RNA interaction coefficients highlighted the specificity preference of p-variants for their expected optimal r-variant binding partner. In most cases, p-variants preferentially edit their cognate r-variants. For example, p1 preferentially edited r1, p2 preferentially edited r2 etc. This is particularly clear for the interactions between m1–m4 and the nucleotides -15 to -12 before the edit site (Fig. 4B). Maximum pooled coefficients for p1 to p30 and r1 to r30 show the RNA base preference of each dsnSc variant in the context of this assay. As we expected based on previous characterisations of the PPR code for PPR motifs (14, 15), Thr in the 5th position of the dsnSc motif strongly favours purines and Asn in the 5th position of the motif strongly favours pyrimidines. The TD combination strongly favours guanine, and the TN combination prefers adenine, but as was reported previously (15), NN and ND motif combinations do not as strongly distinguish between pyrimidines. However, NN shows a slight preference for cytidine (Fig. 4C). Pooled coefficients for interactions between motifs preceding and following an aligned dsnSc motif indicate the identity of neighbouring motifs and nucleotides affects net protein specificity (Fig. 4D).

**Figure 5.**
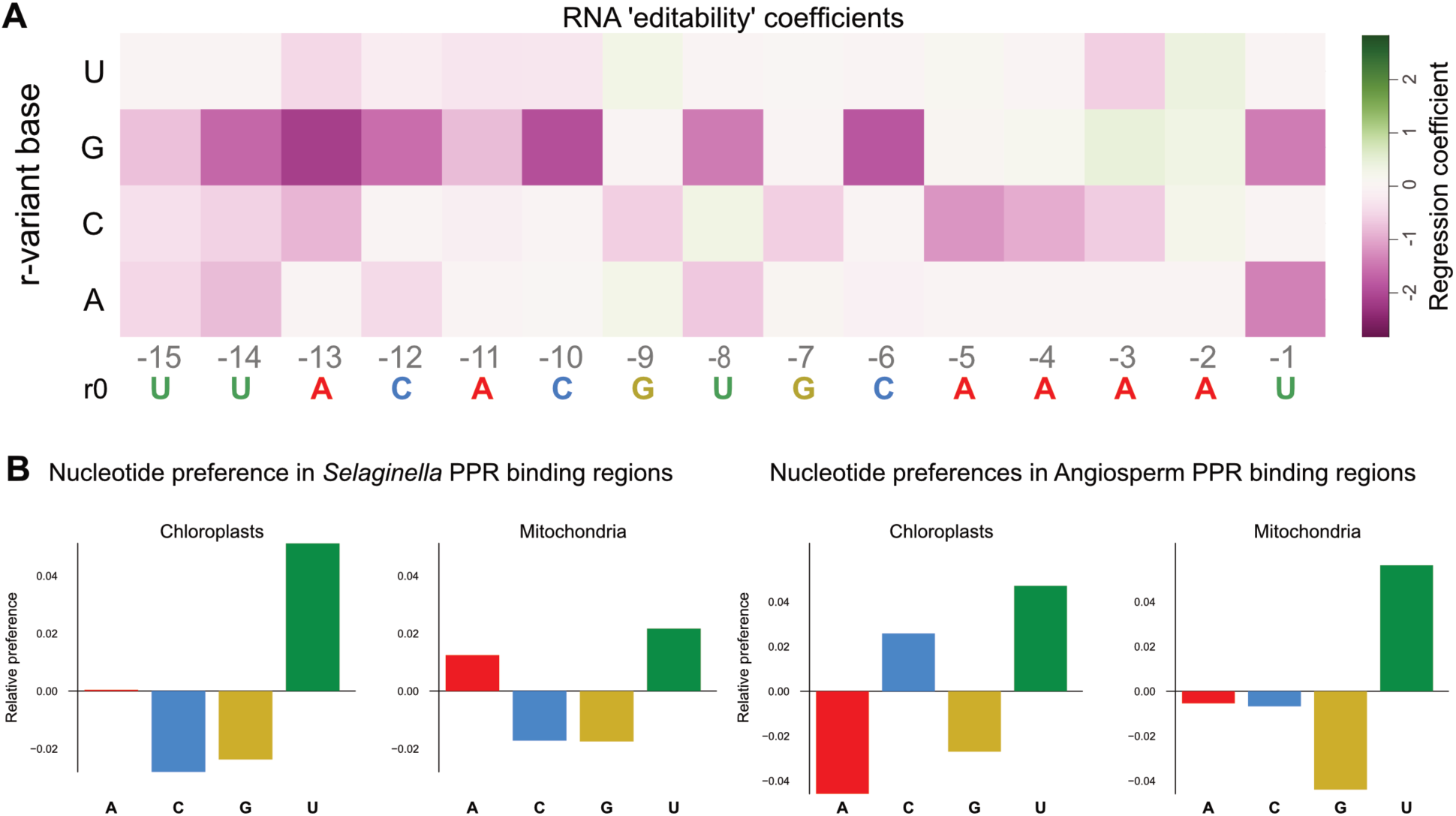
Coefficients for RNA targets show how ‘editable’ RNA variants are relative to the unmodified *rpoA* target. (**A**) RNA coefficients indicating ‘editability’; (**B**) Base composition of PPR protein binding sites inferred from sites -15 to -6 nucleotides upstream of 5213 verified RNA editing sites in *Selaginella kraussiana*, *Selaginella lepidophylla* (54) and *Selaginella uncinata* (55) chloroplasts, and 1410 sites from *Selaginella moellendorffii* mitochondria (66) (left); Base composition of PPR protein binding sites inferred from nucleotides -15 to -6 upstream of 1268 plastid and 7715 mitochondrial RNA editing sites in angiosperms (right).

**Figure 6.**
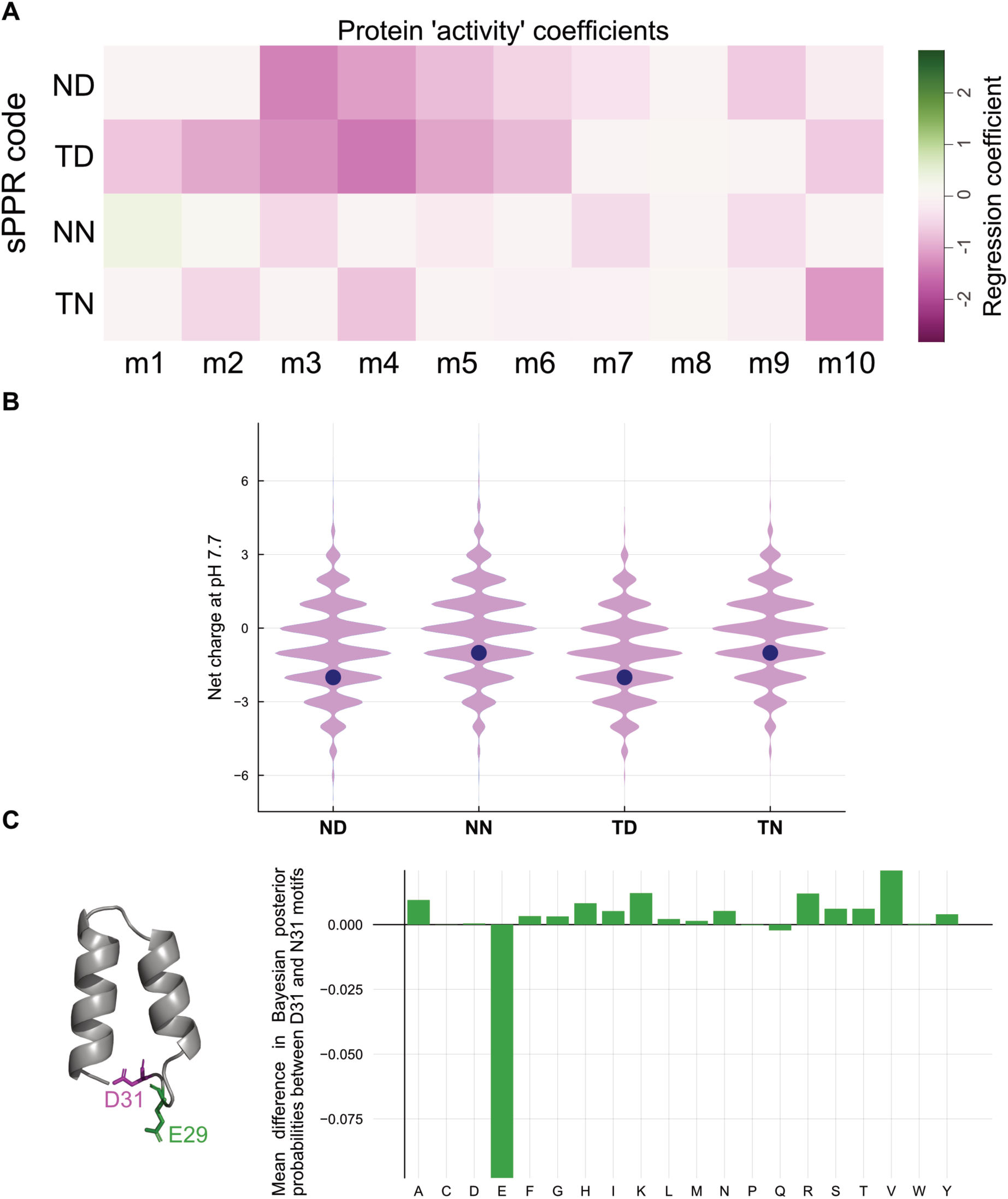
Coefficients for sPPR proteins show protein variant activity relative to unmodified 9SrpoADYW. (**A**) Protein coefficients indicating ‘activity’; (**B**) Distribution of net charge on natural S motifs encoding ND, NN, TD and TN in their 5th and last amino acid specificity combinations. GRASP sPPR motif net charges are indicated as circles; (**C**) D31 affects the probability of E29. The residue in position 29 of the dsnSc motif is a negatively charged Glu residue. The mean difference in Bayesian posterior probabilities of amino acids at position 29 in natural S-class editing proteins shows that Glu is specifically disfavoured in motifs with D31. The dsnSc TD motif was modeled in AlphaFold3 (63) and visualised using PyMol (64).

### Effects of mismatches on RNA editability

RNA coefficients from the regression analysis show the ‘editability’ of each RNA variant, and illustrates the effects of base modification at each position of the RNA target (Fig. 5A). There is a distinct preference for pyrimidines in position -1. Purines in this position resulted in a decrease of editability. Elsewhere in the RNA target, guanine strongly affected target editability. Modification of a base to guanine in positions -15 to -6 resulted in greatly decreased editability. However, in position -3 the r38 A to G modification increased target editability. Conversely, when a nucleotide is modified from guanine to adenine or uridine, for example at positions -7 and -9, editability increases except when changed to a cytidine. Modification of the adenines in positions -5 to -3 to cytidine also reduced editability, but modification of adenine at the -2 position to any other nucleotide increased editability (Fig. 5A).

We hypothesised that guanine may be avoided in the target sequences of natural PPR protein sequences. We extracted nucleotide sequences -4 to -15 positions upstream of RNA editing sites from the chloroplasts and mitochondria of three *Selaginella* lycophytes (54, 55), from which a large proportion of the dsnSc consensus sequence was derived (27). In *Selaginella* chloroplasts and mitochondria cytidine and guanine are disfavoured, adenine favourability varies between species and organelles, and uridine is the most favoured nucleotide (Fig. 5B). This trend is also apparent in angiosperm chloroplasts and mitochondria where RNA editing is dominantly catalysed by PLS PPR proteins. In angiosperm chloroplasts, adenine and guanine are disfavoured and cytidine and uridine are most represented. In angiosperm mitochondria, adenine is slightly disfavoured, cytidine and guanine are disfavored, and uridine is the most represented nucleotide (Fig. 5B).

### Effects of mismatches on sPPR activity

Protein coefficients indicate relative protein editing activity (Fig. 6A). Generally, introducing a mismatch between a motif and its aligned base decreased protein activity, but there was a stronger decrease in variants with an Asp residue in the last amino acid of the modified motif (ND and TD). The effects of Asn in the last position of the dsnSc motif were less pronounced (NN and TN) (Fig. 6A). Modification of the m8 motif appeared to have no effect on protein activity, leading us to investigate whether this was a sign of influence of the preceding and following motifs. Pooled interaction coefficients of motifs with their cognate target showed that TD motifs decrease specific interactions in the preceding and following motifs (Fig. 4D). Interactions between sPPR motifs may be a function of the net charge along the PPR repeat tract. We calculated the net charge of natural S-motifs and compared them to the GRASP dsnSc motifs (Fig 6B). ND and TD motifs are more negatively charged than NN and TN motifs due to the negatively charged Asp residue at position 31 of ND and TD motifs. We hypothesised that the extra negatively charged D31 residue in ND and TD motifs may disrupt neighbouring motifs and contribute to an unfavourable net negative charge in GRASP sPPR proteins. All dsnSc motifs encode a negatively charged Glu residue at position 29 (Fig. 6C). This E29 residue is dominant in natural TN and NN motifs, but when we calculated the frequency of E29 in natural S-class TD and ND motifs, we determined that E29 is specifically disfavoured in motifs with D31 (Fig. 6C).

## Discussion

### GRASP provides a foundation for optimising synthetic RNA binding proteins

The GRASP kit is a set of 42 plasmids which can be combined to assemble synthetic PPR proteins to target 9-, 14-or 19-nucleotide long RNA sequences. Experimental evidence suggests that the optimal length of a PPR motif tract to maximise affinity for a specific RNA target is 11 motifs, and increasing the number of motifs makes designer PPR proteins more prone to off-target RNA binding (67). With the inclusion of an RNA editing domain which includes the P2-L2- and S2 motifs and the E1-E2-DYW domain, a 9 motif sPPR protein assembled with GRASP targets a 15-nucleotide sequence, with specific motif:RNA interactions forming between 10 motifs (9 dsnSc motifs and the P2-motif). The S2-motif was designed to target all four bases, and the L2-, E1- and E2 motifs generally do not strongly influence RNA binding (14, 68, 69). Therefore, the recommended length for an sPPR RNA editing factor is 9 dsnSc motifs. The inclusion of additional plasmids to build 14 and 19 motifs is included for added flexibility and might be desired for certain applications where additional C-terminal motifs will not be added. For example, a 14 motif sPPR may be more effective for applications that require RNA binding, but no enzymatic domain and a 19 motif sPPR protein allows for the generation of higher molecular weight sPPR proteins that are more amenable to structural studies.

GRASP circumvents challenges associated with highly repetitive sequence assembly for PPR proteins and provides a foundational system for integrating synthetic PPR proteins into modular cloning standards. Through reconstruction of a synthetic RNA editing factor and comprehensive modulation of its RNA binding preference against a set of on- and off-target sequences, we demonstrated the use case of both assembling and optimising synthetic PPR RNA editing factors.

### A high-throughput activity assay for synthetic PPR RNA editing proteins

To pair with high-throughput assembly of sPPR proteins, we developed a high-throughput protein activity assay utilising bacterial cell-free protein expression lysate, *in vitro* transcription of stabilised RNA targets, and custom indexation paired with DNA sequencing to rapidly profile RNA editing activity (Supplementary Fig. S4). This assay, although rapid and well suited to high-throughput protein activity profiling, may not fully reflect protein activity *in vivo*. Previous studies of PPR RNA editing *in vitro* required high concentrations of protein to induce RNA editing (23). sPPR protein activity in *in vitro* assays (23, 70), cell-free expression lysates, and in heterologous *in vivo* systems such as in *E. coli* (9, 24, 27) likely requires high protein concentrations as RNA binding is diffusion-dependent, and sPPR proteins are operating outside of their natural context. In plant chloroplasts there is evidence for formation of phase separated granules containing RNA binding proteins interacting with RNA editing cofactor proteins (71, 72) which could create a localised environment of high protein and RNA concentration. Accessory co-factor proteins such as multiple organellar RNA editing factor (MORF) proteins (73, 74) and organelle RNA recognition motif containing (ORRM) proteins (75) increase PPR binding affinity and RNA editing efficiency (24, 76) in angiosperm chloroplasts. While MORF proteins are absent in non-seed plants like lycophytes (1) and are not required for RNA editing by S-type PPR proteins (27), S-type sPPR proteins may still benefit from RNA granule formation and/or interactions with other cofactor proteins *in planta*.

In our high-throughput activity assay, we can control the concentration of *in vitro* transcribed RNA targets incubated with sPPR proteins, but it is difficult to control the concentration of sPPR proteins expressed in cell-free expression lysate. Further confounding factors of protein activity could be the presence of tRNAs, residual RNA molecules in the cell-free expression lysate, and target RNA secondary structures. We estimated the concentration of sPPR proteins in a 1:20 dilution of cell-free expression reactions to be approximately 50 nanomolar. Each expression reaction was incubated with 75 picomolar pooled *in vitro* transcribed RNA. This ratio of 50 nM protein to 75 pM RNA catalysed a maximum C-to-U conversion of ∼23%. Increasing relative protein concentration would increase maximum C-to-U conversion, but as protein concentration increases, target specificity decreases. This high-throughput assay is well suited for rapid assessment of sPPR protein functionality and is sensitive enough to determine differences in sPPR specificity for a controllable, restricted sequence space.

### Effects of motif and target mismatches on off-target activity

The assay of 31 9S_rpoA_DYW protein variants provided a large dataset of RNA editing activity for sPPR proteins targeting variants of a single RNA target, and allowed us to interrogate the effects of single nucleotide variants on sPPR specificity. Every protein variant was active and able to catalyse the conversion of cytidine to uridine in multiple RNA targets (Fig. 4A). Regression coefficients for protein:RNA interactions (Fig. 4B) shows that each variant preferentially edited its cognate target over the 45 off-target sequences present in the expression reaction. However, this effect was most apparent in variants of motifs m1 to m4. Variants of motifs m5 to m9 only slightly preferred their cognate target over off-targets, indicating a greater tolerance to mismatches in these central motifs of the sPPR proteins. The greater importance of N-terminal/5ʹ interactions has been observed in previous studies of synthetic PPR proteins (67, 70). In four designer PPR10 variants described in (67), bind-n-seq experiments showed enrichment for RNAs matching the N-terminus of the protein, with mismatches increasingly tolerated towards the C-terminus. In a separate molecular dynamics study of a synthetic PPR10 protein (70) similarly noted more stable interactions between proteins and RNA targets that matched at the N-terminal/5ʹ end of the complex. The interaction coefficients in 9S_rpoA_DYW m1 to m4 variants supports the hypothesis proposed in (67, 70) that target recognition in synthetic PPR proteins initiates by interaction of N-terminal PPR motifs and cognate RNA sequences, and then the rest of the protein ‘zips-up’ on the RNA. However, observations on natural PPR RNA editing factors are not always in agreement. Mutations of CLB19 C-terminal motifs decreased target specificity, while mutations in N-terminal motifs were more tolerated (68). This may be explained by the differing structure of PLS-class PPR proteins such as CLB19, and their requirement for cofactor proteins such as MORF9 (73, 74) for optimal binding.

RNA coefficients (Fig. 5A) show how ‘editable’ the RNA target is by all protein variants relative to r0. Some effects were expected, such as the inhibitory effect of purines in the -1 position of RNA targets, where there is typically a preference for pyrimidines in natural editing sites (9, 26, 29, 31). For reasons which are unclear, modifications of nucleotides in positions -3 and -2, which align with E1 and E2 motifs, respectively, induced large increases in editability. The E1 and E2 motifs are strongly conserved 34 amino acid motifs that precede the DYW editing domain and are implicated in correctly orienting the DYW domain to its target C (77). E-motifs have been shown to participate in RNA binding, but have a different RNA binding code to P1, L1, S1 and S-type motifs (77). We still know too little about the role of E motifs in RNA editing. While U is generally favourable for RNA editability, modifications in positions -15 to -6 to G strongly decreased editability. Modification of an RNA base from G to A or U (but not C) in positions -9 and -7 increased editability. Generally, it seems that guanine is disfavoured in RNA targets. This could be an artefact of G in the RNA target influencing RNA secondary structure, but RNA target folding predictions do not suggest a major shift in secondary structure, and the decrease in RNA editability is most pronounced with G in positions -15 to -6 where RNA binding specificity is primarily determined. We found that guanine is systematically disfavoured in this region of natural editing sites in *Selaginella* chloroplasts (whilst U is systematically favoured). These preferences may be more general for PPR editing factors, not just S-class proteins as guanine is also disfavoured in angiosperm editing sites (and again uridine is favoured) (Fig. 5B).

Protein coefficients show the activity of each variant relative to p0 (Fig. 6A). The most apparent trend in the motifs was the higher performance of NN and TN motifs than ND and TD motifs. Each dsnSc motif carries a net negative charge, but ND and TD carry one extra negatively charged amino acid residue at D31 (Fig. 6B). Given the decrease in protein activity of ND and TD motif variants, we hypothesised that D31 causes an electrostatic potential imbalance, resulting in the observed decrease in protein activity for D31 motifs (Fig. 6A). Natural S-motifs encoding TD carry a lower net charge than natural NN and TN S-motifs. Therefore, the charge at other positions in the motif must compensate for the presence of D31. The dsnSc motif encodes a negatively charged Glu residue in position 29. Comparison of natural D31 and N31 motifs showed that there is a higher probability of a difference in the frequency of E29 between D31 and N31 motifs than in the frequency of any other amino acids (Fig. 6C). Neutralisation of E29 in dsnSc ND and TD motifs by modification to a neutral amino acid residue may increase protein activity in future GRASP PPR proteins. Modification of motifs generally resulted in a decrease of protein activity, except for the m8 motif which is surrounded by TD motifs. Modification of m8 to any other code had no effect on protein activity (Fig. 6A). This suggests that TD motifs (and sPPR motifs in general) strongly influence flanking motif:base interactions. Motif:base interactions preceding and following a TD:G interaction were specifically reduced (Fig. 4D), suggesting intramolecular forces between motifs affect protein activity. A similar pattern in motif modifications altering flanking sequence preference was observed in off-target editing events in human cell cytosol expressing *Physcomitrium patens* PPR56 (29). Mutation of an S1 TD motif to TN in *Pp*PPR56 altered the preference of the motif from G to A, but also altered the preference of preceding nucleotides from A to C, and the preference of the following nucleotides to strongly prefer G. Mutation of a different S1 motif from TN to TD predictably altered the aligned base from A to G, and altered the preference of the preceding nucleotides from C to U, while the following base preference was unchanged (29). The effects of inter-motif interactions should be considered carefully when choosing targets for sPPR proteins, and may inform the likelihood of off-target RNA editing events.

### Optimisation of consensus sPPR motifs

The GRASP kit uses a PPR code of just four motif-base interactions: ND to target uridine, NN to target cytidine, TN to target adenine and TD to target guanine (Fig. 1B). These fifth and last amino acid combinations were selected for their strong and specific interactions with their cognate base, but natural PPR motifs encode many more fifth and last amino acid combinations which vary in cognate base recognition affinity (15). There are likely many other residues in natural PPR motifs that modulate motif affinity, or form intramolecular interactions with cumulative effects across the PPR motif array as a whole. The dsnSc was derived from the consensus of 3026 S-motifs from 37 land plant species (27) and the motif sequence apart from the fifth and last amino acid are the same in every repeat. The on- and off-target activity data provided by the GRASP high-throughput activity assay has identified several areas which can be targeted for iterative improvement of the GRASP kit.

The dsnSc motif carries a net negative charge, with D31 motifs carrying one extra negatively charged amino acid side chain. With the neutralisation of the charge at E29 in D31 motifs, all dsnSc motifs would carry the same net negative charge. Another position for optimisation is at Q13 of the dsnSc motif. Crystal structures of synthetic PPR proteins showed that the phosphate group of the RNA sugar-phosphate backbone is oriented towards a positive electrostatic region in the first helix of the PPR motif. Salt bridge formation between K13 or R13 in PPR motifs and the RNA phosphate group is essential for RNA binding in some synthetic PPR proteins (78, 79). Substitution of Lys13 to Alanine in the synthetic PPR protein “dPPRU_8_N_2_“ (78), or Lys12 (Lys13 in redefined motif delineation) to Ser12/Glu12 in “cPPR” (18) abolished RNA binding. The dsnSc motif encodes a polar, uncharged Q residue at position 13 which is not capable of salt bridge formation. Modification of Q13 to K13 in tandem with modification of E29 to a neutral residue in ND and TD motifs would completely neutralise the negative charge on dsnSc motifs and may increase sPPR protein activity.

Specific modifications or variants of the E1-E2-DYW domain may also increase the efficacy of sPPR RNA editing factors. In plant chloroplasts and mitochondria, a highly conserved class of editing events creates start codons by converting an ACG codon to an ATG start codon, allowing translation to initiate. As the DYW domain used in 9S_rpoA_DYW is derived from a consensus of thousands of natural DYW domains, it exhibits a preference for pyrimidines reflective of the frequency of pyrimidines in the -1 position relative to natural RNA editing sites. There are likely DYW sequence variants which favour A in position -1 in natural editing factors targeting these sites. Generating consensus sequences based on domains with known targets may increase editing activity for specific target sequences. Alternatively, the use of highly active natural E1-E2-DYW RNA editing domains, such as *Pp*PPR56, could substitute consensus editing domains.

### Perspectives for synthetic PPRs

The ability to rapidly design and assemble novel synthetic PPR proteins to target any desired RNA sequence opens a new avenue of synthetic biology. In nature, PPR proteins are involved in several essential steps in chloroplast and mitochondrial transcript processing. Using the GRASP kit, it is possible to design sPPR proteins to mimic the natural processes of transcript stabilisation, cleavage, splicing and editing. Synthetic PPR proteins fill a gap in chloroplast and mitochondrial synthetic biology capability. Manipulation of chloroplast and mitochondrial genomes is difficult, and in many organisms is not yet possible. The CRISPR revolution remains confined to the nuclear genome due to the difficulty of transporting guide RNAs across organelle membranes. Systems that take advantage of peptide-based nucleic acid binding mechanisms are needed for chloroplast and mitochondrial biotechnology. Methods such as TALENs can be used in place of CRISPR-Cas9 to efficiently edit organelle DNA (43, 44, 80), and PPR proteins can be used in place of CRISPR-Cas13 to edit organelle RNA. An added benefit of sPPR proteins is the ability to implement non-permanent and reversible modifications to RNA, since the modifications introduced are dependent on the expression of the sPPR protein which can be controlled at the nuclear level, or exogenously. sPPR proteins have been shown to work in multiple biological contexts including in bacteria (9, 24, 27, 28), plants, (20, 22, 24, 25) and in human cell culture (28, 29). Though they show great promise for chloroplast and mitochondrial biotechnology, sPPR proteins are not confined to organelles and have been used to edit transcripts in the cytosol (26, 29). Beyond fundamental research, sPPR proteins have the potential to be developed for applications such as stabilising co-factors for mRNA, therapeutics targeting pyrimidine mutations associated with mitochondrial disease, or as molecular switches to control translation.

## Data availability

Plasmid maps of GRASP level -1 modules, level 0 vectors used to assemble 9S*_rpoA_*DYW and variants, and level 1 vectors for 9S*_rpoA_*DYW and variants are provided as GenBank files in the GRASP GitHub repository. The GRASP kit will be made available via Addgene. The code used for analysis of native RNA editing sites and GenBank files are available at https://github.com/farleykvdg/GRASP and is archived at 10.5281/zenodo.15771370.

## Supplementary Data

Supplementary Data are available at *NAR* Online.

## Author contributions

Conceptualisation: F.M.K. and I.S. Investigation: M.D., S.L, A.V., C.C., C.S.B., A.P. and F.M.K. Methodology: F.M.K, A.V., M.D. and I.S. Formal Analysis: I.S. Data curation: L.C. and I.S. Software: I.S. Funding acquisition: I.S. and C.S.B. Supervision: I.S. and C.S.B. Visualisation: F.M.K and I.S. Writing – original draft: F.M.K. Writing – review and editing: M.D., S.Y, A.V., A. P., C.C., L.C., C.S.B, I.S. and F.M.K.

## Funding

This work was supported by the Australian Research Council [grant DP200102981 to I.S. and C.S.B]; and the Australian Research Council Centre of Excellence in Plants for Space [CE230100015].

## Conflict of interest

The authors declare no competing interests.

## Notes

https://github.com/farleykvdg/GRASP

### Competing Interest Statement

The authors have declared no competing interest.

### Summary of Updates

Added author affiliations for 3 - ARC CoE Plants for Space to F.K.V, A.V., L.C., I.S. Added Zenodo DOI to data availability statement.

## References

1. Gutmann, B., Royan, S., Schallenberg-Rüdinger, M., Lenz, H., Castleden, I.R., McDowell, R., Vacher, M.A., Tonti-Filippini, J., Bond, C.S., Knoop, V., et al. (2020) The expansion and diversification of pentatricopeptide repeat RNA-editing factors in plants. Mol. Plant, 13, 215–230.

2. Small, I.D. and Peeters, N. (2000) The PPR motif - a TPR-related motif prevalent in plant organellar proteins. Trends Biochem. Sci., 25, 46–47.

3. Prikryl, J., Rojas, M., Schuster, G. and Barkan, A. (2011) Mechanism of RNA stabilization and translational activation by a pentatricopeptide repeat protein. Proc. Natl. Acad. Sci. U. S. A., 108, 415–420.

4. Zhou, W., Lu, Q., Li, Q., Wang, L., Ding, S., Zhang, A., Wen, X., Zhang, L. and Lu, C. (2017) PPR-SMR protein SOT1 has RNA endonuclease activity. Proc. Natl. Acad. Sci. U. S. A., 114, E1554–E1563.

5. Huynh, S.D., Melonek, J., Colas des Francs-Small, C., Bond, C.S. and Small, I. (2023) A unique C-terminal domain contributes to the molecular function of Restorer-of-fertility proteins in plant mitochondria. New Phytol., 240, 830–845.

6. Chateigner-Boutin, A.-L., des Francs-Small, C.C., Delannoy, E., Kahlau, S., Tanz, S.K., de Longevialle, A.F., Fujii, S. and Small, I. (2011) OTP70 is a pentatricopeptide repeat protein of the E subgroup involved in splicing of the plastid transcript rpoC1: OTP70 is required for splicing of rpoC1. Plant J., 65, 532–542.

7. de Longevialle, A.F., Meyer, E.H., Andrés, C., Taylor, N.L., Lurin, C., Millar, A.H. and Small, I.D. (2007) The pentatricopeptide repeat gene OTP43 is required for trans-splicing of the mitochondrial nad1 Intron 1 in Arabidopsis thaliana. Plant Cell, 19, 3256–3265.

8. Lurin, C., Andrés, C., Aubourg, S., Bellaoui, M., Bitton, F., Bruyère, C., Caboche, M., Debast, C., Gualberto, J., Hoffmann, B., et al. (2004) Genome-wide analysis of Arabidopsis pentatricopeptide repeat proteins reveals their essential role in organelle biogenesis. Plant Cell, 16, 2089–2103.

9. Oldenkott, B., Yang, Y., Lesch, E., Knoop, V. and Schallenberg-Rüdinger, M. (2019) Plant-type pentatricopeptide repeat proteins with a DYW domain drive C-to-U RNA editing in *Escherichia coli*. *Commun*. Biol., 2, 85.

10. Boussardon, C., Avon, A., Kindgren, P., Bond, C.S., Challenor, M., Lurin, C. and Small, I. (2014) The cytidine deaminase signature HxE(x)n CxxC of DYW1 binds zinc and is necessary for RNA editing of ndhD-1. New Phytol., 203, 1090–1095.

11. Salone, V., Rüdinger, M., Polsakiewicz, M., Hoffmann, B., Groth-Malonek, M., Szurek, B., Small, I., Knoop, V. and Lurin, C. (2007) A hypothesis on the identification of the editing enzyme in plant organelles. FEBS Lett., 581, 4132–4138.

12. Takenaka, M., Takenaka, S., Barthel, T., Frink, B., Haag, S., Verbitskiy, D., Oldenkott, B., Schallenberg-Rüdinger, M., Feiler, C.G., Weiss, M.S., et al. (2021) DYW domain structures imply an unusual regulation principle in plant organellar RNA editing catalysis. Nat. Catal., 4, 510–522.

13. Fujii, S., Bond, C.S. and Small, I.D. (2011) Selection patterns on restorer-like genes reveal a conflict between nuclear and mitochondrial genomes throughout angiosperm evolution. Proc. Natl. Acad. Sci. U. S. A., 108, 1723–1728.

14. Barkan, A., Rojas, M., Fujii, S., Yap, A., Chong, Y.S., Bond, C.S. and Small, I. (2012) A combinatorial amino acid code for RNA recognition by pentatricopeptide repeat proteins. PLoS Genet., 8, e1002910.

15. Yan, J., Yao, Y., Hong, S., Yang, Y., Shen, C., Zhang, Q., Zhang, D., Zou, T. and Yin, P. (2019) Delineation of pentatricopeptide repeat codes for target RNA prediction. Nucleic Acids Res., 47, 3728–3738.

16. Yang, F., Vincis Pereira Sanglard, L., Lee, C.-P., Ströher, E., Singh, S., Oh, G.G.K., Millar, A.H., Small, I. and Colas des Francs-Small, C. (2022) Knockdown of mitochondrial*atp1*mRNA by a custom-designed pentatricopeptide repeat protein alters F1FoATP synthase. bioRxiv, 10.1101/2022.11.08.515711.

17. Colas des Francs-Small, C., Vincis Pereira Sanglard, L. and Small, I. (2018) Targeted cleavage of nad6 mRNA induced by a modified pentatricopeptide repeat protein in plant mitochondria. *Commun*. Biol., 1, 166.

18. Coquille, S., Filipovska, A., Chia, T., Rajappa, L., Lingford, J.P., Razif, M.F.M., Thore, S. and Rackham, O. (2014) An artificial PPR scaffold for programmable RNA recognition. Nat. Commun., 5, 5729.

19. Gully, B.S., Shah, K.R., Lee, M., Shearston, K., Smith, N.M., Sadowska, A., Blythe, A.J., Bernath-Levin, K., Stanley, W.A., Small, I.D., et al. (2015) The design and structural characterization of a synthetic pentatricopeptide repeat protein. Acta Crystallogr. D Biol. Crystallogr., 71, 196–208.

20. Mathieu, S., Lesch, E., Garcia, S., Stéfanie, G., Schallenberg-Rüdinger, M. and Hammani, K. (2024) *De novo*RNA base editing in plant organelles with engineered synthetic P-type PPR editing factors. bioRxiv, 10.1101/2024.09.13.612905.

21. Kwok van der Giezen, F., Honkanen, S., Colas des Francs-Small, C., Bond, C. and Small, I. (2024) Applications of synthetic pentatricopeptide repeat proteins. Plant Cell Physiol., 65, 503–515.

22. Manavski, N., Mathieu, S., Rojas, M., Méteignier, L.-V., Brachmann, A., Barkan, A. and Hammani, K. (2021) In vivo stabilization of endogenous chloroplast RNAs by customized artificial pentatricopeptide repeat proteins. Nucleic Acids Res., 49, 5985–5997.

23. Hayes, M.L. and Santibanez, P.I. (2020) A plant pentatricopeptide repeat protein with a DYW-deaminase domain is sufficient for catalyzing C-to-U RNA editing in vitro. J. Biol. Chem., 295, 3497–3505.

24. Royan, S., Gutmann, B., Colas des Francs-Small, C., Honkanen, S., Schmidberger, J., Soet, A., Sun, Y.K., Vincis Pereira Sanglard, L., Bond, C.S. and Small, I. (2021) A synthetic RNA editing factor edits its target site in chloroplasts and bacteria. *Commun*. Biol., 4, 545.

25. Manavski, N., Abdel-Salam, E., Schwenkert, S., Kunz, H.-H., Brachmann, A., Leister, D. and Meurer, J. (2025) Targeted introduction of premature stop codon in plant mitochondrial mRNA by a designer pentatricopeptide repeat protein with C-to-U editing function. Plant J., 121, e17247.

26. Thielen, M., Gärtner, B., Knoop, V., Schallenberg-Rüdinger, M. and Lesch, E. (2024) Conquering new grounds: plant organellar C-to-U RNA editing factors can be functional in the plant cytosol. Plant J., 119, 895–915.

27. Bernath-Levin, K., Schmidberger, J., Honkanen, S., Gutmann, B., Sun, Y.K., Pullakhandam, A., Colas des Francs-Small, C., Bond, C.S. and Small, I. (2021) Cofactor-independent RNA editing by a synthetic S-type PPR protein. Synth. Biol., 7, ysab034.

28. Ichinose, M., Kawabata, M., Akaiwa, Y., Shimajiri, Y., Nakamura, I., Tamai, T., Nakamura, T., Yagi, Y. and Gutmann, B. (2022) U-to-C RNA editing by synthetic PPR-DYW proteins in bacteria and human culture cells. *Commun*. Biol., 5, 968.

29. Lesch, E., Schilling, M.T., Brenner, S., Yang, Y., Gruss, O.J., Knoop, V. and Schallenberg-Rüdinger, M. (2022) Plant mitochondrial RNA editing factors can perform targeted C-to-U editing of nuclear transcripts in human cells. Nucleic Acids Res., 50, 9966–9983.

30. Gerke, P., Szövényi, P., Neubauer, A., Lenz, H., Gutmann, B., McDowell, R., Small, I., Schallenberg-Rüdinger, M. and Knoop, V. (2020) Towards a plant model for enigmatic U-to-C RNA editing: the organelle genomes, transcriptomes, editomes and candidate RNA editing factors in the hornwort *Anthoceros agrestis*. New Phytol., 225, 1974–1992.

31. Kwok van der Giezen, F.M., Viljoen, A., Campbell-Clause, L., Dao, N.T., Colas des Francs-Small, C. and Small, I. (2024) Insights into U-to-C RNA editing from the lycophyte *Phylloglossum drummondii*. Plant J., 119, 445–459.

32. Hayes, M.L., Garcia, E.T., Chun, S.O. and Selke, M. (2024) Crosslinking of base-modified RNAs by synthetic DYW-KP base editors implicates an enzymatic lysine as the nitrogen donor for U-to-C RNA editing. J. Biol. Chem., 300, 107454.

33. Ichinose, M., Teramoto, T., Nakamura, I., Shimajiri, Y., Yagi, Y. and Gutmann, B. (2025) Fine-tuning of the PPR protein directs the RNA editing activity toward C-to-U or U-to-C conversion. Sci. Rep., 15, 6288.

34. Abudayyeh, O.O., Gootenberg, J.S., Franklin, B., Koob, J., Kellner, M.J., Ladha, A., Joung, J., Kirchgatterer, P., Cox, D.B.T. and Zhang, F. (2019) A cytosine deaminase for programmable single-base RNA editing. Science, 365, 382–386.

35. Kavuri, N.R., Ramasamy, M., Qi, Y. and Mandadi, K. (2022) Applications of CRISPR/Cas13-based RNA editing in plants. Cells, 11, 2665.

36. Abudayyeh, O.O., Gootenberg, J.S., Essletzbichler, P., Han, S., Joung, J., Belanto, J.J., Verdine, V., Cox, D.B.T., Kellner, M.J., Regev, A., et al. (2017) RNA targeting with CRISPR-Cas13. Nature, 550, 280–284.

37. Engler, C., Kandzia, R. and Marillonnet, S. (2008) A one pot, one step, precision cloning method with high throughput capability. PLoS One, 3, e3647.

38. Weber, E., Engler, C., Gruetzner, R., Werner, S. and Marillonnet, S. (2011) A modular cloning system for standardized assembly of multigene constructs. PLoS One, 6, e16765.

39. Engler, C., Youles, M., Gruetzner, R., Ehnert, T.-M., Werner, S., Jones, J.D.G., Patron, N.J. and Marillonnet, S. (2014) A golden gate modular cloning toolbox for plants. ACS Synth. Biol., 3, 839–843.

40. Moore, S.J., Lai, H.-E., Kelwick, R.J.R., Chee, S.M., Bell, D.J., Polizzi, K.M. and Freemont, P.S. (2016) EcoFlex: A multifunctional MoClo kit for E. coli synthetic biology. ACS Synth. Biol., 5, 1059–1069.

41. Iverson, S.V., Haddock, T.L., Beal, J. and Densmore, D.M. (2016) CIDAR MoClo: Improved MoClo assembly standard and new E. coli part library enable rapid combinatorial design for synthetic and traditional biology. ACS Synth. Biol., 5, 99–103.

42. Zhang, S., Wang, J. and Wang, J. (2020) One-day TALEN assembly protocol and a dual-tagging system for genome editing. ACS Omega, 5, 19702–19714.

43. Arimura, S.-I. (2022) MitoTALENs: A method for targeted gene disruption in plant mitochondrial genomes. Methods Mol. Biol., 2363, 335–340.

44. Nakazato, I. and Arimura, S.-I. (2025) Targeted C-to-T base editing in the Arabidopsis Plastid genome. Curr. Protoc., 5, e70075.

45. Kang, B.-C., Bae, S.-J., Lee, S., Lee, J.S., Kim, A., Lee, H., Baek, G., Seo, H., Kim, J. and Kim, J.-S. (2021) Chloroplast and mitochondrial DNA editing in plants. Nat. Plants, 7, 899–905.

46. Abil, Z., Denard, C.A. and Zhao, H. (2014) Modular assembly of designer PUF proteins for specific post-transcriptional regulation of endogenous RNA. J. Biol. Eng., 8, 7.

47. Yagi, Y., Teramoto, T., Kaieda, S., Imai, T., Sasaki, T., Yagi, M., Maekawa, N. and Nakamura, T. (2022) Construction of a versatile, programmable RNA-binding protein using designer PPR proteins and its application for splicing control in mammalian cells. Cells, 11, 3529.

48. Cheng, S., Gutmann, B., Zhong, X., Ye, Y., Fisher, M.F., Bai, F., Castleden, I., Song, Y., Song, B., Huang, J., et al. (2016) Redefining the structural motifs that determine RNA binding and RNA editing by pentatricopeptide repeat proteins in land plants. Plant J., 85, 532–547.

49. Olins, P.O., Devine, C.S., Rangwala, S.H. and Kavka, K.S. (1988) The T7 phage gene 10 leader RNA, a ribosome-binding site that dramatically enhances the expression of foreign genes in Escherichia coli. Gene, 73, 227–235.

50. Kluesner, M.G., Tasakis, R.N., Lerner, T., Arnold, A., Wüst, S., Binder, M., Webber, B.R., Moriarity, B.S. and Pecori, R. (2021) MultiEditR: The first tool for the detection and quantification of RNA editing from Sanger sequencing demonstrates comparable fidelity to RNA-seq. Mol. Ther. Nucleic Acids, 25, 515–523.

51. Blind, M., Kolanus, W. and Famulok, M. (1999) Cytoplasmic RNA modulators of an inside-out signal-transduction cascade. Proc. Natl. Acad. Sci. U. S. A., 96, 3606–3610.

52. Hunsicker, A., Steber, M., Mayer, G., Meitert, J., Klotzsche, M., Blind, M., Hillen, W., Berens, C. and Suess, B. (2009) An RNA aptamer that induces transcription. Chem. Biol., 16, 173–180.

53. Bushnell, B., Rood, J. and Singer, E. (2017) BBMerge -Accurate paired shotgun read merging via overlap. PLoS One, 12, e0185056.

54. Smith, D.R. (2020) Unparalleled variation in RNA editing among Selaginella plastomes. Plant Physiol., 182, 12–14.

55. Oldenkott, B., Yamaguchi, K., Tsuji-Tsukinoki, S., Knie, N. and Knoop, V. (2014) Chloroplast RNA editing going extreme: more than 3400 events of C-to-U editing in the chloroplast transcriptome of the lycophyte *Selaginella uncinata*. RNA, 20, 1499–1506.

56. One Thousand Plant Transcriptomes Initiative (2019) One thousand plant transcriptomes and the phylogenomics of green plants. Nature, 574, 679–685.

57. Beick, S., Schmitz-Linneweber, C., Williams-Carrier, R., Jensen, B. and Barkan, A. (2008) The pentatricopeptide repeat protein PPR5 stabilizes a specific tRNA precursor in maize chloroplasts. Mol. Cell. Biol., 28, 5337–5347.

58. Pfalz, J., Bayraktar, O.A., Prikryl, J. and Barkan, A. (2009) Site-specific binding of a PPR protein defines and stabilizes 5’ and 3’ mRNA termini in chloroplasts. EMBO J., 28, 2042– 2052.

59. Haïli, N., Arnal, N., Quadrado, M., Amiar, S., Tcherkez, G., Dahan, J., Briozzo, P., Colas des Francs-Small, C., Vrielynck, N. and Mireau, H. (2013) The pentatricopeptide repeat MTSF1 protein stabilizes the nad4 mRNA in Arabidopsis mitochondria. Nucleic Acids Res., 41, 6650–6663.

60. Potapov, V., Ong, J.L., Kucera, R.B., Langhorst, B.W., Bilotti, K., Pryor, J.M., Cantor, E.J., Canton, B., Knight, T.F., Evans, T.C., Jr, et al. (2018) Comprehensive profiling of four base overhang ligation fidelity by T4 DNA ligase and application to DNA assembly. ACS Synth. Biol., 7, 2665–2674.

61. Guillaumot, D., Lopez-Obando, M., Baudry, K., Avon, A., Rigaill, G., Falcon de Longevialle, A., Broche, B., Takenaka, M., Berthomé, R., De Jaeger, G., et al. (2017) Two interacting PPR proteins are major Arabidopsis editing factors in plastid and mitochondria. Proc. Natl. Acad. Sci. U. S. A., 114, 8877–8882.

62. Lorenz, R., Bernhart, S.H., Höner Zu Siederdissen, C., Tafer, H., Flamm, C., Stadler, P.F. and Hofacker, I.L. (2011) ViennaRNA Package 2.0. Algorithms Mol. Biol., 6, 26.

63. Abramson, J., Adler, J., Dunger, J., Evans, R., Green, T., Pritzel, A., Ronneberger, O., Willmore, L., Ballard, A.J., Bambrick, J., et al. (2024) Accurate structure prediction of biomolecular interactions with AlphaFold 3. Nature, 630, 493–500.

64. Schrödinger, LLC (2015) The PyMOL Molecular Graphics System, Version 1.8.

65. Zuker, M. (2003) Mfold web server for nucleic acid folding and hybridization prediction. Nucleic Acids Res., 31, 3406–3415.

66. Hecht, J., Grewe, F. and Knoop, V. (2011) Extreme RNA editing in coding islands and abundant microsatellites in repeat sequences of *Selaginella moellendorffii* mitochondria: the root of frequent plant mtDNA recombination in early tracheophytes. Genome Biol. Evol., 3, 344–358.

67. Miranda, R.G., McDermott, J.J. and Barkan, A. (2018) RNA-binding specificity landscapes of designer pentatricopeptide repeat proteins elucidate principles of PPR-RNA interactions. Nucleic Acids Res., 46, 2613–2623.

68. Kindgren, P., Yap, A., Bond, C.S. and Small, I. (2015) Predictable alteration of sequence recognition by RNA editing factors from Arabidopsis. Plant Cell, 27, 403–416.

69. Matsuda, T., Sugita, M. and Ichinose, M. (2020) The L motifs of two moss pentatricopeptide repeat proteins are involved in RNA editing but predominantly not in RNA recognition. PLoS One, 15, e0232366.

70. Marzano, N., Johnston, B., Paudel, B.P., Schmidberger, J., Jergic, S., Böcking, T., Agostino, M., Small, I., van Oijen, A.M. and Bond, C.S. (2024) Single-molecule visualization of sequence-specific RNA binding by a designer PPR protein. Nucleic Acids Res., 52, 14154–14170.

71. Legen, J., Lenzen, B., Kachariya, N., Feltgen, S., Gao, Y., Mergenthal, S., Weber, W., Klotzsch, E., Zoschke, R., Sattler, M., et al. (2024) A prion-like domain is required for phase separation and chloroplast RNA processing during cold acclimation in Arabidopsis. Plant Cell, 36, 2851–2872.

72. Chodasiewicz, M., Sokolowska, E.M., Nelson-Dittrich, A.C., Masiuk, A., Beltran, J.C.M., Nelson, A.D.L. and Skirycz, A. (2020) Identification and characterization of the heat-induced plastidial stress granules reveal new insight into Arabidopsis stress response. Front. Plant Sci., 11, 595792.

73. Yan, J., Zhang, Q., Guan, Z., Wang, Q., Li, L., Ruan, F., Lin, R., Zou, T. and Yin, P. (2017) MORF9 increases the RNA-binding activity of PLS-type pentatricopeptide repeat protein in plastid RNA editing. Nat. Plants, 3, 17037.

74. Takenaka, M., Zehrmann, A., Verbitskiy, D., Kugelmann, M., Härtel, B. and Brennicke, A. (2012) Multiple organellar RNA editing factor (MORF) family proteins are required for RNA editing in mitochondria and plastids of plants. Proc. Natl. Acad. Sci. U. S. A., 109, 5104–5109.

75. Shi, X., Bentolila, S. and Hanson, M.R. (2016) Organelle RNA recognition motif-containing (ORRM) proteins are plastid and mitochondrial editing factors in Arabidopsis. Plant Signal. Behav., 11, e1167299.

76. Lombana, J.M., Hanson, M.R. and Bentolila, S. (2025) Deciphering the role of accessory proteins in Arabidopsis chloroplast editosomes via interaction with a synthetic PPR-PLS factor in E. coli. Nucleic Acids Res., 53.

77. Ruwe, H., Gutmann, B., Schmitz-Linneweber, C., Small, I. and Kindgren, P. (2019) The E domain of CRR2 participates in sequence-specific recognition of RNA in plastids. New Phytol., 222, 218–229.

78. Shen, C., Zhang, D., Guan, Z., Liu, Y., Yang, Z., Yang, Y., Wang, X., Wang, Q., Zhang, Q., Fan, S., et al. (2016) Structural basis for specific single-stranded RNA recognition by designer pentatricopeptide repeat proteins. Nat. Commun., 7, 11285.

79. Yin, P., Li, Q., Yan, C., Liu, Y., Liu, J., Yu, F., Wang, Z., Long, J., He, J., Wang, H.-W., et al. (2013) Structural basis for the modular recognition of single-stranded RNA by PPR proteins. Nature, 504, 168–171.

80. Arimura, S.-I. and Nakazato, I. (2024) Genome editing of plant mitochondrial and chloroplast genomes. Plant Cell Physiol., 65, 477–483.

