## Supplementary Figure Legends for "GRASP: a modular toolkit for building synthetic pentatricopeptide repeat RNA-binding proteins" (from another version of this preprint: version 1, 2025-06-30)

**Supplementary Figure S1.** Catalogue of GRASP plasmids. (**A**) Extension of a 9-motif GRASP sPPR is achieved through addition of 5 or 10 dsnSc motifs in the middle of the PPR motif array (orange and blue motifs in 14S and 19S AlphaFold models) (63). To assemble a 14-motif protein, parts 1A-B-C-D-14E, 14A-B-C-D-1E, 2A-B-C-D-2E are assembled into a level 1 plasmid construct. To assemble a 19-motif protein, parts 1A-B-C-D-14E, 14A-B-C-D-19E, 19A-B-C-D-1E, 2A-B-C-D-2E are assembled into a level 1 plasmid construct. Compatible assembly junctions are shown. (**B**) The 42 plasmids that make up the GRASP kit are comprised of nucleotide randomised variants of parts 1A, B, C, D, 1E, 2A, 2E, 14A, 14E, 19A and 19E which differ in the amino acid identity of the 5th and last residues which confer specificity for an RNA base. Assembly junction variants of 1A are also included. (AGGT) may be used when fusing the PPR array to an N-terminal sequence, and (AATG) may be used when translation initiation is intended to start at the first PPR motif. (**C**) Choosing the appropriate parts for a given RNA sequence is achieved through matching the PPR code nomenclature to its RNA base. For example, in the given 9 nucleotide sequence of CGUACAGUA, parts 1A_N and B_NT combine to reconstitute a dsnSc motif with the code NN, which preferentially binds cytosine. Parts B_NT and C_DN reconstitute a dsnSc motif with the code TD, which preferentially binds guanidine, and so on. A script to simplify part selection according to the desired RNA target is provided.

**Supplementary Figure S2.** Assembly, expression and functional assay of 19S*_rpoA_*DYW. (**A**) An sPPR protein encoding 19 motifs targeting a co-transcribed, extended *rpoA* sequence was assembled with the GRASP kit. (**B**) 19S*_rpoA_*DYW was expressed in *E. coli* and purified using nickel immobilised metal affinity chromatography. The 19S*_rpoA_*DYW protein is visible in protein fractions at the estimated molecular weight of 118 kDa. (**C**) Sanger sequencing of the co-transcribed RNA editing target site showed that 19S*_rpoA_*DYW is functional and capable of RNA editing. Sanger chromatogram ratios were measured with MultiEditR (50).

**Supplementary Figure S3.** Uncropped SDS-PAGE and Western blot of 9S*_rpoA_*DYW expressed in *E. coli* cell-free expression lysate. (**A**) Acrylamide gel showing expression of 9S*_rpoA_*DYW in triplicate run alongside the BioRad unstained protein standards molecular weight marker, a no template control and the NEBExpress cell-free expression positive control plasmid DHFR. (**B**) Western blot of the acrylamide gel detecting 9S*_rpoA_*DYW with an anti-6XHistidine tag secondary antibody.

**Supplementary Figure S4.** Overview of the high-throughput activity assay utilising bacterial cell-free expression lysate. Plasmid constructs encoding sPPRs are expressed for 2.5 hours at 37℃. In tandem, DNA oligonucleotides are transcribed *in vitro* to generate stabilised RNA editing targets. Pooled RNA targets are added to protein expression reactions a further 2.5 hours at 37℃, and then are reverse transcribed, pooled, and made into an DNA sequencing library and sequenced on an Illumina MiSeq.
