## Supplementary Table Legends for "GRASP: a modular toolkit for building synthetic pentatricopeptide repeat RNA-binding proteins" (from another version of this preprint: version 1, 2025-06-30)

**Table S1.** GRASP level -1 module plasmids.

**Table S2.** Variant 9S*_rpoA_*DYW sPPR proteins and their cognate RNA targets.

**Table S3.** Variant *rpoA* RNA targets flanked by custom NGS primer binding regions and 5ʹ and 3ʹ stem loop sequences.

**Table S4.** List of primers used.

**Table S5.** RNA stem-loop target fold predictions generated by the Mfold web server RNA4.0 at 37℃ Linear Max 1 folding (65).
